# Inherited input and local transformations shape the spatiotemporal organization of pathway specific striatal signals for motivated behavior

**DOI:** 10.64898/2026.06.04.730000

**Authors:** Zicheng Zhang, Yizhuo Ding, Mai-Anh Vu, Lydia Mroz, Yuxin Tong, Mark Howe

## Abstract

Adaptive behavior requires neural circuits to link sensory events with their location, predictive value, and temporal relationship to outcomes. Although striatal circuits are implicated in this process, it remains unclear how motivationally relevant signals are organized across striatal regions and direct- and indirect-pathway spiny projection neurons, and which components reflect afferent input versus local transformations. Using striatum-wide calcium recordings during visual conditioning in mice, we found that learned cue value, reward proximity, cue location, and lick-related behavior were encoded in distinct regions and time windows. These signals included both pathway-convergent representations and pathway-opponent dynamics. Striatum-wide measurements of glutamatergic input targeted to each SPN subtype revealed that rapid cue-location and lick-related signals were present in afferent input to both pathways, consistent with inherited representations. In contrast, pathway-opponent pDMS value signals and dSPN-selective pVLS ramping were absent from corresponding glutamatergic input dynamics, indicating local striatal transformations. These findings reveal region-specific input-output transformations that organize striatal signals for distinct components of motivated behavior.

## Introduction

Learned cues guide behavior by signaling where relevant events occur, what outcomes they predict, and when those outcomes are likely to arrive. Adaptive behavior requires neural circuits to extract these distinct features and convert them into output signals that influence orienting, action selection, and anticipatory responses. The striatum is a major site where sensory and associative signals can be converted into motivationally relevant output. Striatal spiny projection neurons (SPNs) receive convergent glutamatergic input from cortical and subcortical circuits ^1–3^, and the postsynaptic influence of these inputs is shaped by neuromodulatory signals, including dopamine and acetylcholine, that encode features of reward prediction, salience, and movement state^4–9^. Perturbation studies have linked region-specific striatal function to aspects of cue-outcome learning, cue-driven approach, incentive motivation, and anticipatory responding in Pavlovian and instrumental contexts^10–14^. Electrophysiological and optical recordings have identified region-specific striatal responses to conditioned cues, including signals related to cue identity, cue-outcome associations, expected value, and motivational significance^15–20^. Striatal populations also show anticipatory or ramping dynamics during delays and approach to reward^21–24^. These findings support the view that striatal circuits participate in transforming cue-related information into signals that guide learning and motivated behavior. However, many structures that project to striatum are also implicated in these processes and contain neurons that signal features similar to those observed in SPNs^25–30^. Thus, SPN activity may reflect either the inheritance of computations generated upstream or local transformations performed within striatal circuits. Distinguishing these alternatives is essential for identifying what computations are implemented by striatal output neurons.

A key substrate for local transformation is the division of striatal projection neurons into direct-and indirect-pathway SPN subtypes (dSPNs and iSPNs), which project through distinct basal ganglia output pathways ^6,31^. Both populations receive topographically organized glutamatergic input from overlapping sources, but they differ in the modulatory and local circuit mechanisms that shape how these inputs influence output^2,3,6,32,33^. dSPNs and iSPNs express largely distinct dopamine receptor classes, allowing dopamine to differentially modulate excitatory input and synaptic plasticity across the two pathways^6,9^. They also participate in asymmetric lateral inhibition and receive distinct modulation from local GABAergic and cholinergic interneurons^34–36^. These features provide mechanisms through which a shared or partially overlapping glutamatergic input scaffold could be strengthened, weakened, or filtered differently across pathways. Consistent with this possibility, prior in vivo studies have reported distinct or opposing dSPN and iSPN representations of stimulus valence, movement, or motivational salience, and some of this divergence is mirrored in downstream basal ganglia output structures^19,20,37–40^. Other studies, however, report largely similar cue, value, and movement-related signals across pathways^20,41–44^. These findings suggest that divergent dSPN and iSPN encoding is not a uniform circuit property, but may depend on the variable being encoded, the timing of the signal, and the striatal region in which it is measured.

The anatomical organization of the striatum is likely central to this problem. Striatal territories receive topographically organized afferent input and participate in largely parallel basal ganglia-thalamocortical circuits^2,3,45^. This regional organization is preserved in downstream basal ganglia output pathways, creating distinct input-output architectures across striatal territories^46,47^. Large-scale recordings have revealed functionally specialized striatal dynamics across striatal territories, and perturbation studies indicate that these territories make separable contributions to cue-guided learning and behavior^12,13,48–50^. These observations indicate that the relationship between afferent input and SPN dynamics is regionally organized. In some territories, SPN dynamics may closely mirror local inputs, whereas in others, striatal circuit mechanisms may transform regionally distinct input signals into pathway-specific dSPN and iSPN dynamics.

Testing whether striatal signals are inherited from afferent input or locally transformed within specific SPN pathways has been limited by methodological constraints. Electrophysiological recordings can sample broadly across striatal territories, but cell-type identification across large populations generally requires optotagging or other low-throughput approaches, limiting region-wide comparisons of dSPN and iSPN dynamics. Calcium imaging and fiber photometry allow pathway-specific recordings, but most prior studies have focused on single regions and have not directly compared SPN calcium dynamics with local glutamatergic input. As a result, it remains unclear how pathway-specific representations of sensory and motivational variables are organized across the striatum, and which components are inherited from afferents versus locally transformed within striatal circuits.

Here, we measured striatum-wide calcium and glutamate dynamics from dSPNs and iSPNs during Pavlovian visual conditioning in mice. These recordings reveal that dSPN and iSPN calcium dynamics exhibit divergent encoding of learned cue-value and reward proximity within particular striatal regions, and this divergence is not present in corresponding glutamatergic input dynamics, indicative of local, cell-type specific transformations. For task-related representations in other regions, encoding in calcium dynamics largely mirrored that in the input signals, indicating inheritance from upstream regions. These findings provide a framework for understanding how input scaffolds are transformed across striatal circuits to support distinct aspects of motivated behavior and learning.

## Results

### Striatum-wide, pathway-specific population Ca^2+^ recordings reveal spatially and temporally organized cue responses during Pavlovian conditioning

We used a Pavlovian delay conditioning task designed to isolate striatal representations of stimulus features relevant for motivated behavior, including cue identity, cue location, relative value, and temporal proximity to reward (Fig. 1A). Head-fixed mice were presented on each trial with either a reward-associated visual cue (CS+) or a non-reward-associated cue (CS-), followed by water delivery after a 3 s delay on CS+ trials only. In a subset of mice, cues were presented pseudorandomly at one of three positions spanning the visual field: contralateral, central, or ipsilateral to the implanted hemisphere. Mice were positioned on a spherical treadmill, allowing spontaneous two-dimensional locomotion, although locomotion had no effect on task outcome. Across training, mice acquired robust cue discrimination, developing significantly greater anticipatory licking during the delay period on CS+ trials than on CS- trials (Fig. 1B-D). This discrimination was present across all cue locations, indicating that mice learned the cue-reward association independently of stimulus position (Supplementary Fig. 1A).

**Figure 1.**
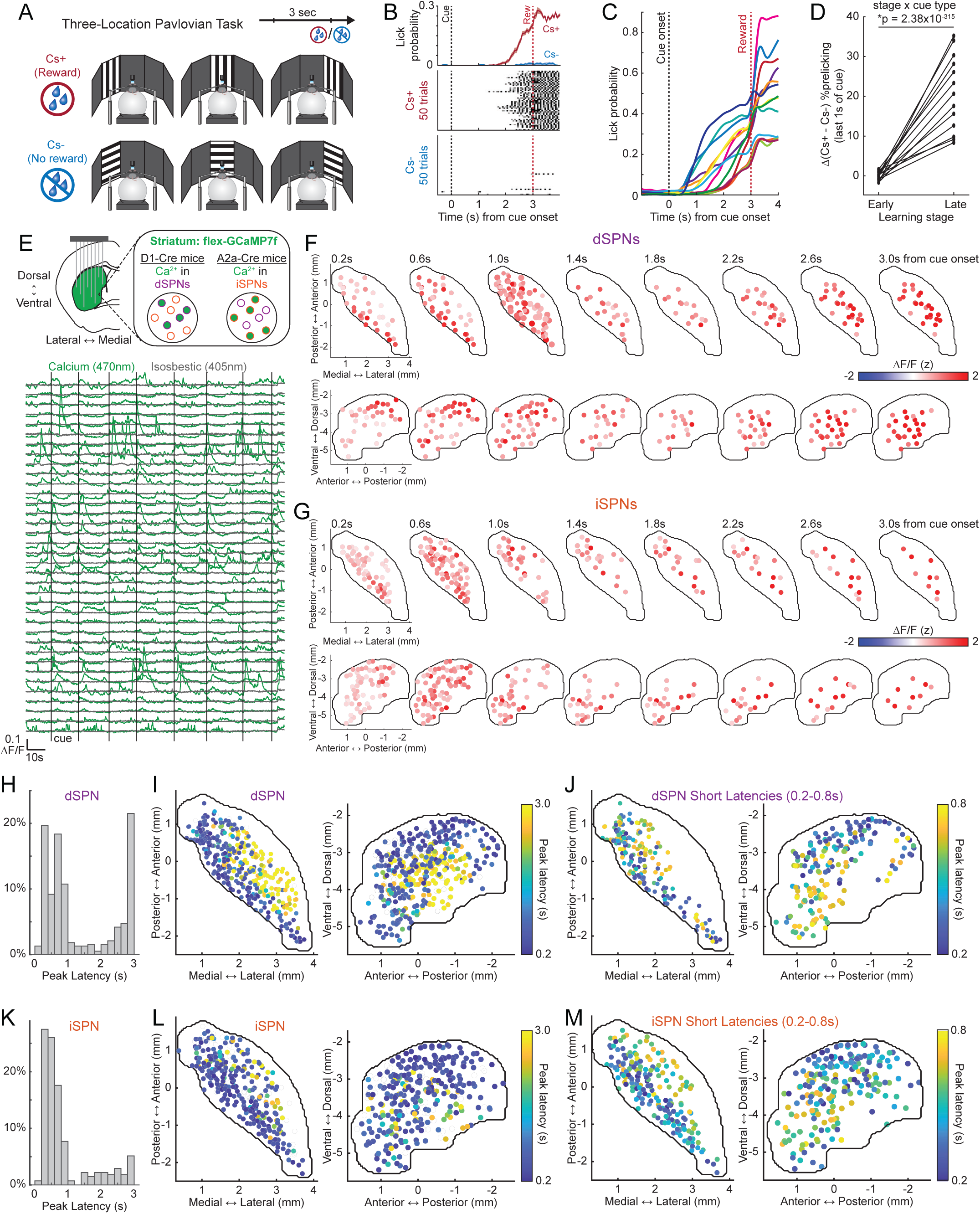
Striatum-wide calcium recordings reveal spatially organized cue responses during Pavlovian conditioning (A) Schematic of the three-location Pavlovian task. On each trial, mice were presented with a reward-predictive visual cue (CS+) or non-rewarded visual cue (CS-) at one of three screen locations, followed by reward delivery 3 s after cue onset on CS+ trials. (B) Representative licking behavior from one mouse aligned to cue onset. Top, mean lick probability on CS+ and CS- trials. Shaded regions indicate SEM. Bottom, lick rasters for CS+ and CS- trials; each row is one trial and each tick is one lick. Black dashed line indicates cue onset and red dashed line indicates reward delivery. (C) Mean lick probability on CS+ trials across individual mice. Each colored line represents one mouse. Black dashed line indicates cue onset and red dashed line indicates reward delivery. (D) Cue discrimination increased across learning. Each line represents one mouse and connects early and late learning values. Cue discrimination was quantified as the CS+ minus CS- lick probability during the final second before reward delivery. Significance was assessed using a linear mixed-effects model of lick probability as a function of learning stage, cue type, and their interaction. Stage × cue type interaction, p = 2.38 × 10^-315^; n = 13 mice. (E) Top, schematic of the implantation and calcium-sensor targeting strategy used to record dSPN and iSPN activity across the striatum. jGCaMP7f expression was restricted to dSPNs or iSPNs in separate D1-Cre and A2A-Cre cohorts. Bottom, representative simultaneously recorded dSPN Ca^2+^ fluorescence signals across sites during a single session from one D1-Cre mouse. Each row corresponds to one recording site. Green traces show calcium-dependent 470 nm jGCaMP7f signals and gray traces show 405 nm isosbestic control signals. Dashed vertical lines indicate cue presentations. (F) Spatial maps from an example D1-Cre mouse showing learned-stage CS+ cue-evoked dSPN activity across successive 0.4 s bins from cue onset. A total of 48 recording sites were included in the analysis. Each circle represents a site with a significant cue response in the corresponding time bin, and color indicates its mean cue-evoked activity. Top row, horizontal view; bottom row, sagittal view. (G) Same as (F), for an example A2a-Cre mouse with 93 recording sites included. (H) Distribution of peak response latencies across CS+ cue-responsive dSPN sites. Peak latency was measured within the 0.2-3.0 s post-cue window. (I) Horizontal and sagittal striatal maps of peak response latencies for CS+ cue-responsive dSPN sites. Each circle indicates one recording site and is colored by the latency of its dominant cue-evoked peak. Unfilled circles indicate recording sites that did not show a significant cue response at any time point. (J) Same as (I), restricted to short-latency dSPN sites with peak responses between 0.2 and 0.8 s after cue onset. (K) Same as (H), for iSPN sites. (L) Same as (I), for iSPN sites. (M) Same as (J), for short-latency iSPN sites. For (H)-(M), analyses include CS+ cue-responsive sites from the learned phase: dSPNs, n = 333 sites from 7 mice; iSPNs, n = 280 sites from 6 mice.

To measure pathway-specific striatal population dynamics, we recorded Ca^2+^ signals from 34-81 sites per mouse distributed throughout the striatal volume using targeted arrays of small-diameter optical fibers^51^ (46 μm; Fig. 1E). Expression of the green fluorescent Ca^2+^ indicator jGCaMP7f was restricted to dSPNs or iSPNs using separate cohorts of D1-Cre and A2A-Cre mice, respectively (n = 8 D1-Cre mice, 430 fibers; n = 6 A2A-Cre mice, 329 fibers). Fiber locations were identified post mortem using micro-CT reconstruction, and spatial coverage was comparable across dSPN and iSPN cohorts (Supplementary Fig. 1B,C). Each fiber sampled fluorescence from a small, non-overlapping tissue volume, enabling parallel measurements of local population activity across the striatum^51^. Cue-evoked Ca^2+^ transients varied across recording sites and were absent in quasi-simultaneous 405 nm isosbestic control recordings, ruling out substantial motion or hemodynamic contamination (Fig. 1E; see Methods).

After discriminative learning, reward-predictive CS+ cues evoked widespread but spatiotemporally organized Ca^2+^ responses across a large fraction of both dSPN and iSPN recording sites (see Methods, 333/364 dSPN sites from 7 mice; 280/329 iSPN sites from 6 mice; Fig. 1F-M). Many cue-responsive sites peaked within the first second after cue onset (Fig. 1F-I,K,L). Within this short-latency population, the earliest responses were concentrated in pDMS, followed by slightly later responses in anterior ventral striatum, including nucleus accumbens regions (Fig. 1F,G,J,M). Both response components were observed in dSPNs and iSPNs. In addition to these early cue responses, a separate population of sites showed later peaks near the expected time of reward delivery. These late responses were concentrated in posterior ventrolateral striatum and were observed preferentially in dSPNs (Fig. 1F,H,I). Thus, reward-predictive cues recruited multiple temporally distinct response components organized across anatomically segregated striatal territories.

Ca^2+^ responses were also observed during CS- trials, but, on average, these responses peaked at significantly shorter latencies than CS+ responses in both pathways (Supplementary Fig. 1D; Wilcoxon rank-sum test, dSPN: p = 4.87 × 10^-28^; iSPN: p = 9.46 × 10^-10^), and delayed response components were largely absent. Very few significant cue-evoked signals were detected in isosbestic recordings, further confirming that these dynamics did not arise from motion or hemodynamic artifacts (4/157 dSPN sites from 3 mice, 5/93 iSPN sites from 2 mice).

To distinguish learned cue responses from pre-existing visual responses, we next examined activity evoked by the same visual stimuli before conditioning in a subset of mice. Before learning, visual cues evoked short-latency responses in both dSPNs and iSPNs, with no difference in the proportion of visually responsive sites or peak latencies between pathways (Supplementary Fig. 1E,F; difference in % sites: linear mixed effect model, p = 0.177; difference in peak latency: linear mixed effect model, p = 0.518). These responses did not distinguish between the future CS+ and CS- stimuli, with only a small fraction of sites showing any pre-learning difference between cue identities (5/366 dSPN sites and 4/329 iSPN sites). Pre-learning responses were localized primarily to pDMS in both pathways, and the significant response regions overlapped across dSPNs and iSPNs (Supplementary Fig. 1G-I; see Methods, spatial overlap test: p=0.029). Thus, visual stimuli initially evoke rapid, anatomically localized responses that are shared across striatal output pathways, whereas longer-latency cue-locked and delay-period activity emerges after cues acquire reward-predictive value.

### Cue location is encoded by learning invariant, short-latency dSPN and iSPN responses in posterior dorsomedial striatum

To quantify how striatal activity encoded distinct task variables across regions and cell types, we fit linear regression models to .LF/F traces from each recording site (Fig. 2A; Supplementary Fig. 2A). Models included finite impulse response (FIR) kernels for discrete task variables, including cue value and cue location, in addition to continuous behavioral regressors capturing licking and treadmill movement (Methods). We first examined encoding of cue location in the visual field. Location selectivity was quantified using FIR terms corresponding to cue position relative to the implanted hemisphere (Fig. 1A; Fig. 2A; Methods). Individual sites exhibited robust cue-evoked responses with either little location selectivity or strong contralateral-biased location selectivity, illustrating how the model separated general cue responses from location selective components (Fig. 2B,C).

**Figure 2.**
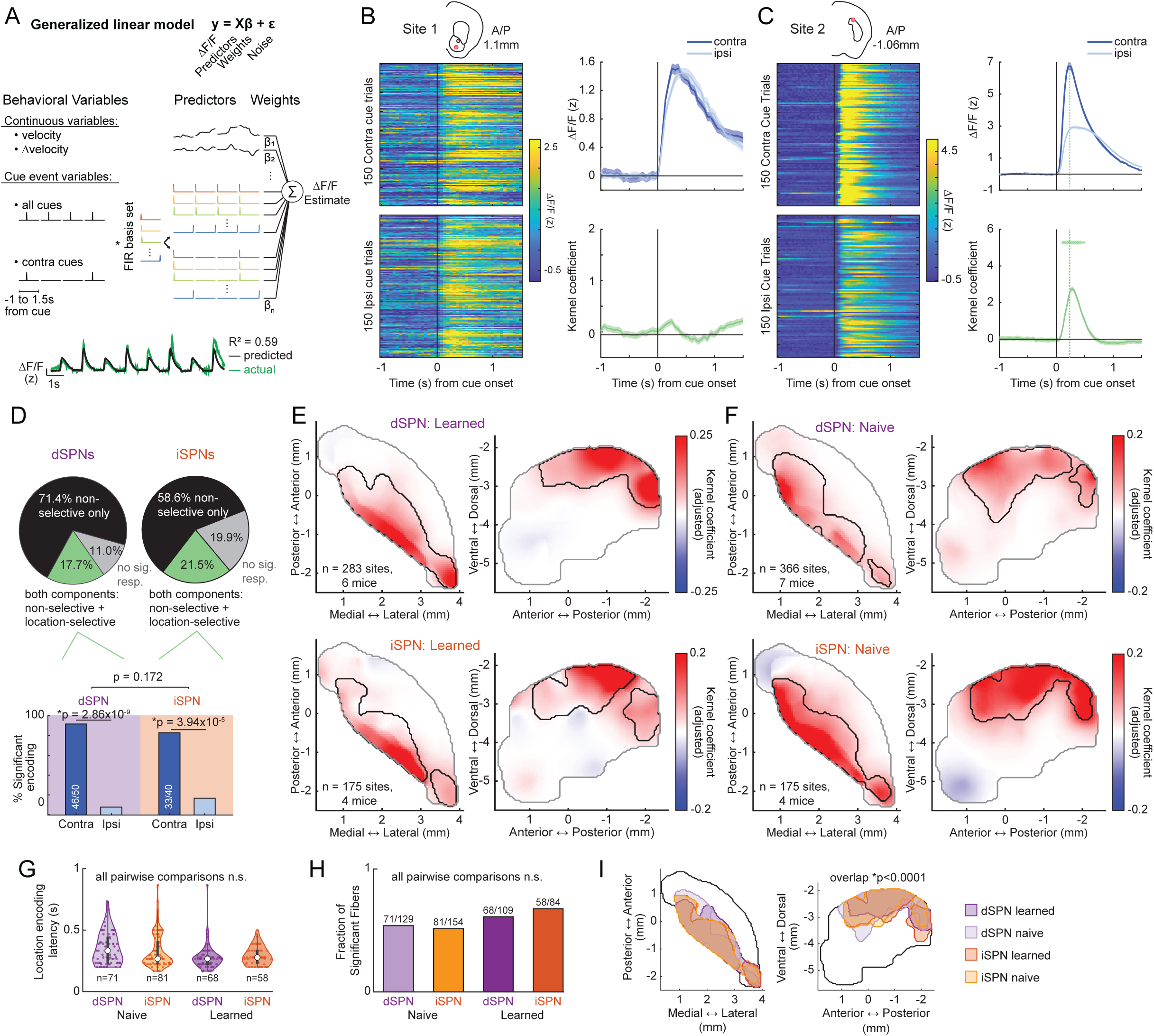
Cue location is encoded by shared short-latency dSPN and iSPN responses in posterior dorsomedial striatum. (A) Generalized linear model capture location-selective cue responses. Continuous predictors included velocity and change in velocity. Cue event predictors included all-cue events and contralateral-cue events, each expanded with a finite impulse response basis spanning -1 to 1.5 s from cue onset. Weighted predictors were summed to estimate the calcium .LF/F signal. Bottom, representative model fit for one recording site; green trace shows measured .LF/F and black trace shows model prediction. (B) Example iSPN recording site without significant location encoding. Left, anatomical location of the site and trial-by-trial responses to contralateral and ipsilateral cue presentations. Each row in the heat maps represents one trial. Right, mean .LF/F responses to contralateral and ipsilateral cues and corresponding model kernels. The green trace shows the location-selective kernel. No significant location encoding was detected in this example. (C) Same as (B), for an example iSPN site with significant contralateral-biased location encoding. Green asterisks mark time points with significant location encoding, and the green vertical line indicates the time at which the location-selective kernel reached 80% of its peak amplitude. Shaded regions indicate SEM. (D) Classification of learned-stage recording sites. Top, pie charts show the fraction of sites with only location-nonselective cue responses, both location-nonselective and location-selective responses, or no significant cue response. The no-sig category includes sites with location encoding but no significant cue response. Bottom, cue-location preference among location-selective sites. Location-selective sites were significantly biased toward contralateral cue locations in both dSPNs and iSPNs, and the fraction of contralateral-preferring sites did not differ between cell types. Chi-square goodness-of-fit tests, dSPN: p = 2.86 × 10^-9^; iSPN: p = 3.94 × 10^-5^; chi-square test of independence, p = 0.172. (E) Smoothed horizontal and sagittal maps of learned-stage location encoding in dSPNs and iSPNs. Voxel color indicates the peak coefficient amplitude of the location-selective kernel after spatial smoothing. Red indicates contralateral preference and blue indicates ipsilateral preference. Black contours mark significant group-level spatial encoding regions. (F) Same as (E), for naive-stage dSPN and iSPN recordings. (G) Distribution of location-encoding latencies within the significant contour across naive and learned-stage dSPN and iSPN datasets. Dots indicate individual recording sites. Pairwise comparisons were not significant (p > 0.05). Linear mixed model with latency as a function of task phase and cell type and mouse as a random effect. (H) Fraction of significant sites within the spatial-encoding region within the significant contour across cell type and learning stage. Numbers above bars indicate significant sites over total sites. The fraction of significant sites did not differ across cell types or task stages. Generalized linear mixed model with significance status predicted by task stage and cell type and mouse as a random effect. (I) Overlay of significant spatial encoding regions from learned and naive dSPN and iSPN datasets. Spatial encoding regions significantly overlapped across all four conditions (p < 0.0001). For (E)-(I), learned-stage datasets include dSPNs, n = 283 sites from 6 mice, and iSPNs, n = 175 sites from 4 mice; naive-stage datasets include dSPNs, n = 366 sites from 7 mice, and iSPNs, n = 175 sites from 4 mice. Spatial overlap was assessed using the spatial overlap test described in Methods. Asterisks in model kernels indicate significant time points.

Although significant cue-evoked Ca^2+^ responses were present at most recording sites after learning (Fig. 1), only a minority displayed significant location selectivity (Fig. 2D). The proportion of ipsi vs contra location-selective sites did not differ between dSPNs and iSPNs (chi-square test of independence, p = 0.172; Fig. 2D). Across both cell types, location-selective sites predominantly responded more strongly to contralateral than ipsilateral cues (chi-square test for goodness of fit, dSPN: p = 2.86 × 10^-9^; iSPN: p = 3.94 × 10^-5^; Fig. 2D). To determine how location encoding was organized across the striatum, we reconstructed three-dimensional contours encompassing regions with significant location-related model coefficients (Methods; Fig. 2E). Spatial consistency across animals was evaluated using bootstrap-based significance tests (Supplementary Fig. 2B-E; Methods). This analysis revealed a clear anatomical concentration of location-selective activity in posterior dorsomedial striatum (pDMS) in both dSPNs and iSPNs (Fig. 2E). In contrast, anterior and ventral striatal regions showed little location selectivity despite exhibiting robust cue-evoked responses after learning (Fig. 1F,G,I,L; Fig. 2E). Thus, cue location was not represented uniformly across all cue-responsive striatal territories, but was selectively concentrated in pDMS.

Because cues presented at different visual locations could evoke different orienting, locomotor, or licking responses, we performed additional analyses to determine whether location selectivity was explained by behavior. Locomotion parameters were included in the model to partially account for movement-related activity, and we repeated the location analysis after matching contralateral and ipsilateral cue trials for linear and angular velocity (Methods; Supplementary Fig. 3A-G). Location-encoding regions were preserved in these movement-matched trials and significantly overlapped with the regions identified in the full dataset (Supplementary Fig. 3G; spatial overlap test: p<0.0001). Regional patterns of location encoding were also preserved when we restricted the analysis to trials without short-latency licking during the cue period (Methods; Supplementary Fig. 3H-J; spatial overlap test: p<0.0001). Together, these analyses indicate that location selectivity primarily reflected encoding of the cue location in the visual field rather than systematic differences in motor output or reward expectation.

Location-selective responses emerged at similarly short latencies in dSPNs and iSPNs (Fig. 2G; linear mixed effect model, dSPN learned vs iSPN learned, p = 0.51), and their anatomical distribution closely matched the short-latency cue responses observed before and after learning (Fig. 1J,M; Supplementary Fig. 1F-I, Fig. 3J). Consistent with this, location encoding was already present prior to conditioning in both pathways (Fig. 2F). The fraction of significant sites within the location-encoding region and the latencies of location-selective responses did not differ across cell type or learning stage (Fig. 2G,H; mixed-effects models for latency and response significance). Cue-location encoding in pDMS was also consistent across individual mice before conditioning (Supplementary Fig. 2F-I), and the significant location-encoding regions overlapped across dSPNs, iSPNs, naive, and learned conditions (Fig. 2I; spatial overlap test: p<0.0001). Together, these findings indicate that visual cue location is encoded by a stable, short-latency pDMS response shared across SPN populations and learning stages, consistent with a pre-existing sensory representation of cue location.

**Figure 3.**
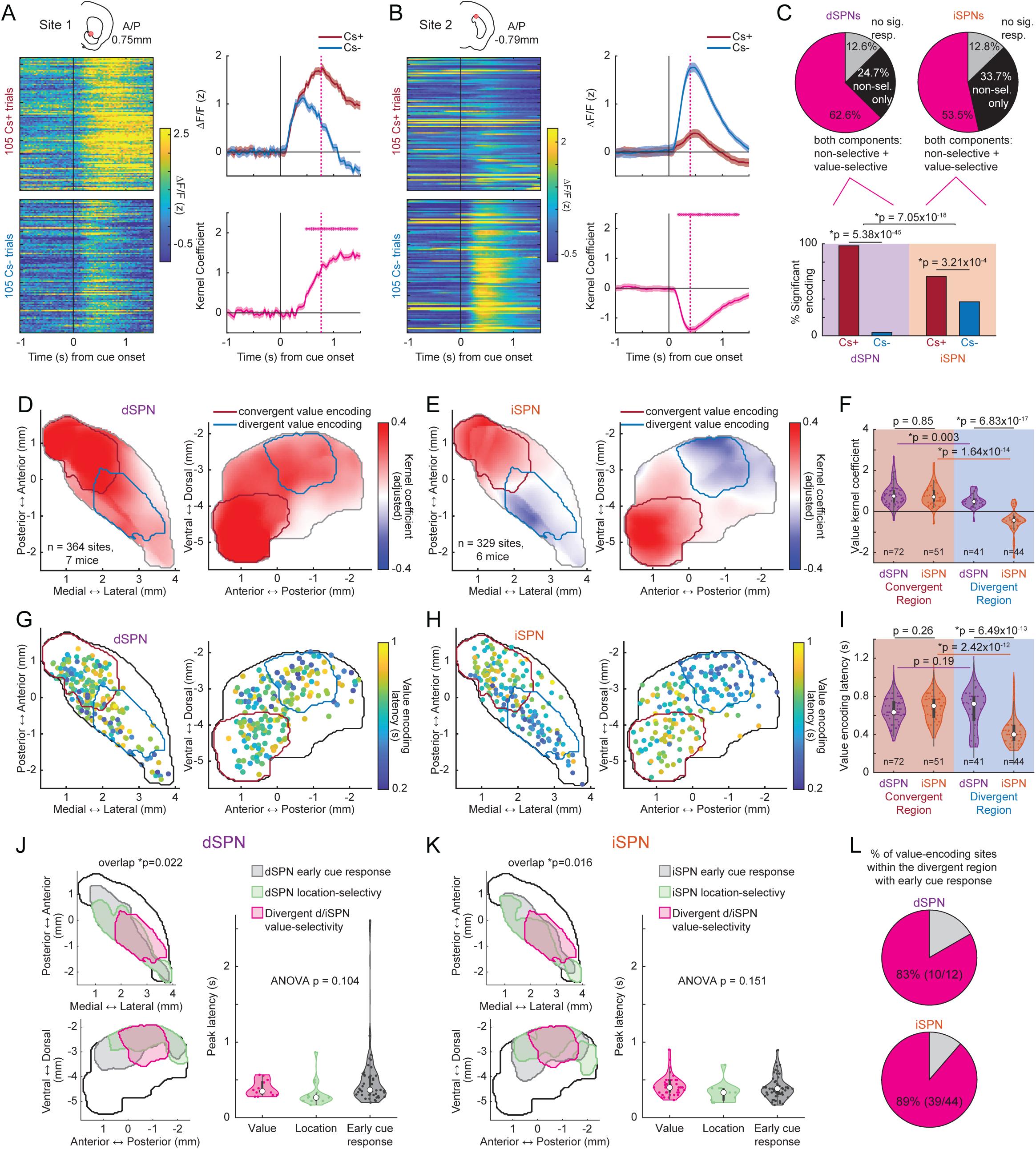
Cue value is encoded by anatomically distinct convergent and pathway-opponent SPN signals. (A) Example learned-stage iSPN recording site with CS+ preference. Left, anatomical location of the site (top) and trial-by-trial responses to 105 CS+ trials and 105 CS- trials. Each row in the heat maps represents one trial. Right, top: mean .LF/F responses to CS+ and CS- cues. Shaded regions indicate SEM. Right, bottom: value-selective kernel. Asterisks mark time points with significant encoding. The magenta vertical line indicates the time at which the value-selective kernel reached 80% of peak amplitude. (B) Same as (A), for a different example iSPN site with CS- preference. (C) Classification of learned-stage recording sites. Top, pie charts show the fraction of sites with only value-nonselective cue responses, both value-selective and value-nonselective cue responses, or sites without a significant cue response for both CS+ and CS-. The no-sig category includes sites with value encoding but no significant cue response. Bottom, cue preference among value-selective sites. Cue preference was significantly non-uniform in both cell types and differed between cell types. Chi-square goodness-of-fit tests, dSPN: p = 5.38 × 10^-45^; iSPN: p = 3.21 × 10^-4^; chi-square test of independence, p = 7.05 × 10 ^-18^. (D) Smoothed horizontal and sagittal maps of learned-stage dSPN value encoding. Voxel color indicates the value-kernel coefficient amplitude after spatial smoothing, with red indicating positive values (CS+ preference) and blue indicating negative values (CS- preference). Red contours denote the significant convergent value-encoding region and blue contours denote the significant divergent value-encoding region. (E) Same as (D), for learned-stage iSPN value encoding. (F) Value-kernel coefficients for dSPN and iSPN sites within the convergent and divergent value-encoding regions. Violin plots show distributions, and dots indicate individual recording sites. Value-kernel amplitudes were similar between dSPN and iSPN sites in the convergent region, but differed between cell types in the divergent region. Linear mixed model with kernel amplitude as a function of region and cell type and mouse as a random effect; convergent dSPN vs iSPN, p = 0.85; divergent dSPN vs iSPN, p = 6.83 × 10^-17^; dSPN convergent vs divergent, p = 0.003; iSPN convergent vs divergent, p = 1.64 × 10 ^-14^. (G) Horizontal and sagittal maps of peak latencies for value-encoding dSPN sites. Each circle indicates one significant value-encoding site and is colored by value-encoding latency. Red and blue contours denote the convergent and divergent value-encoding regions, respectively. (H) Same as (G), for iSPN sites. (I) Value-encoding latencies within the convergent and divergent value-encoding regions. Violin plots show distributions, and dots indicate individual recording sites. iSPN sites in the divergent region encoded value at shorter latency than dSPN sites in the same region and iSPN sites in the convergent region. Linear mixed model with latency as a function of region and cell type and mouse as a random effect; convergent dSPN vs iSPN, p = 0.26; divergent dSPN vs iSPN, p = 6.49 × 10^-13^; dSPN convergent vs divergent, p = 0.19; iSPN convergent vs divergent, p = 2.42 × 10^-12^. (J) Overlay of significant contours for dSPN early cue-response (naive stage, gray), location-selective (green), and divergent value-encoding (magenta) regions in horizontal and sagittal views. The three regions significantly overlapped (p = 0.022). Peak latencies for dSPN value-encoding, location-encoding, and early cue-response sites did not differ (ANOVA, p = 0.104). Dots indicate individual recording sites. (K) Same as (J), for iSPN sites. The three regions significantly overlapped (p = 0.016), and peak latencies did not differ across categories (ANOVA, p = 0.151). (L) Pie charts showing the fraction of dSPN and iSPN value-encoding sites within the divergent region that also exhibited an early cue response in the naive stage. dSPNs, 83% (10/12); iSPNs, 89% (39/44). For (D)-(E), learned-stage maps include dSPNs, n = 364 sites from 7 mice, and iSPNs, n = 329 sites from 6 mice. For (G)-(H), latency maps include value-encoding learned-stage sites. Spatial overlap was assessed using the spatial overlap test described in Methods. Asterisks in model kernels indicate significant time points.

### Anatomically distinct striatal territories encode cue value with shared and pathway-opponent dynamics at different latencies

We next asked whether cue responses encoded learned cue value differently across striatal regions and SPN pathways. We first focused on short latency cue-evoked responses occurring within the first second after cue onset (Fig. 1H,J,K,M). To quantify relative value encoding, we derived GLM kernels at each site that reflected differences between CS+ and CS- responses within this early cue window (Methods; Fig. 3A,B; Supplementary Fig. 4A,B). Individual sites showed either stronger responses to the rewarded CS+ cue or stronger responses to the unrewarded CS- cue, illustrating how the model isolated value selective cue responses (Fig. 3A,B). Value encoding was widespread in both cell-types, but cue preferences differed markedly between dSPNs and iSPNs (Fig. 3C, chi-square test of independence, p = 7.05 × 10^-18^). In dSPNs, value-selective responses were overwhelmingly CS+ preferring (221/229 sites significantly CS+ preferring, chi-square test for goodness of fit, p = 5.38 × 10^-45^; Fig. 3C,D,F; Supplementary Fig. 4C,E). By contrast, iSPNs showed a more heterogeneous distribution, with a significant fraction of sites responding more strongly to CS- than CS+ cues (Fig. 3C,E,F; Supplementary Fig. 4D,F). CS- preferences were observed almost exclusively in iSPNs (65/73 CS- preferring sites) and were present across iSPN mice (Supplementary Fig. 4G). Thus, although both pathways broadly encoded learned cue value, pathway divergence emerged because CS- preferring value signals were selectively expressed in iSPNs.

This pathway divergence was strongly structured anatomically. CS+ preferring dSPN responses were distributed broadly but were strongest and most concentrated in anterior ventral striatum (Fig. 3D). iSPN CS+ preferring responses were concentrated in the same anterior ventral territory, whereas CS- preferring iSPN responses were localized primarily to posterior dorsomedial striatum (pDMS; Fig. 3E). To quantify this regional organization, we defined a continuous three-dimensional striatal volume based on the cell-type differences in value-encoding (Methods). This divergence was localized to the pDMS region (Fig. 3D,E; Methods; center coordinates: AP = -0.65, ML = 2.55, DV = -2.75). A second region was defined in the anterior ventral striatum, where dSPN and iSPN value coding showed strong convergence across pathways, with both populations preferring CS+ (center coordinates: AP = 0.85, ML = 1.45, DV = -4.6). Average value kernels in anterior ventral striatum were strongly positive and did not differ significantly between pathways (linear mixed model of kernel amplitude as a function of region and cell type with random intercept for mouse, p = 0.85; Fig. 3F). In contrast, value kernels in pDMS were significantly divergent and opposite in sign, with positive value coefficients in dSPNs and negative value coefficients in iSPNs (linear mixed model of kernel amplitude as a function of region and cell type with random intercept for mouse, p = 6.83 × 10^-17^; Fig. 3F). These regional differences were consistently present within individual mice, indicating that the regional distribution of cell-type specific cue-value encoding was robust across mice (Supplementary Fig. 4E,F).

Because CS+ and CS- trials could differ in movement or licking profiles, we next tested whether the regional value-encoding patterns were explained by value-related behavioral differences. CS+ and CS- trials showed differences in linear and angular velocity in several mice (Supplementary Fig. 6A,B). We therefore repeated the value-encoding analysis after matching subsets of trials for profiles of cue-evoked linear and angular velocity. Both the convergent anterior ventral value-encoding region and the divergent pDMS region were preserved in velocity matched subsets of trials (Supplementary Fig. 6C), and these regions significantly overlapped with those identified in the full dataset (volume overlap test: convergent region: p<0.0001, divergent region: p=0.006, Supplementary Fig. 6D). Value-encoding regions were also preserved when we restricted the analysis to trials without short-latency prelicking during the delay period (spatial overlap test: convergent region p<0.0001, divergent region p=0.002, Supplementary Fig. 6E). Thus, the regional and pathway-specific value signals were not explained by systematic differences in movement or licking between CS+ and CS- trials.

The two value-coding regions also differed in temporal dynamics. Value-kernel peaks occurred at shorter latencies in the divergent pDMS region than in the convergent anterior ventral region, with the clearest difference observed in iSPNs (linear mixed model of latency as a function of region and cell type with random intercept for mouse, p = 2.42 × 10^-12^; Fig. 3G-I; Supplementary Fig. 4H). In contrast, dSPN value-encoding latencies in pDMS were bimodally distributed, with both short- and longer-latency components (Fig. 3I; Supplementary Fig. 4H,I). We therefore used k-means clustering to separate pDMS dSPN value-encoding sites into fast and slow latency subgroups (Supplementary Fig. 4I). The fast pDMS value signals were anatomically and temporally aligned with the pre-learning sensory and cue-location responses described above. In dSPNs, the fast value-coding region overlapped significantly with both the location-selective region and the pre-learning early cue-response region (spatial overlap test, p = 0.022; Fig. 3J), and the latencies of value, location, and pre-learning cue responses did not differ significantly (ANOVA, p = 0.104; Fig. 3J). Similarly, in iSPNs, the divergent value-coding region overlapped significantly with location-selective and pre-learning cue-response regions (spatial overlap test, p = 0.016; Fig. 3K), and response latencies were indistinguishable across value, location, and pre-learning cue-response categories (ANOVA, p = 0.151; Fig. 3K). Consistent with this spatial and temporal alignment, nearly all value-encoding sites in the divergent pDMS region also showed early cue responses before learning (10/12 dSPN sites and 39/44 iSPN sites; Fig. 3L). By contrast, value coding in anterior ventral striatum emerged at longer latencies, consistent with the delayed cue responses that developed through conditioning (Fig. 1I,L; Fig. 3G-I; Supplementary Fig. 4H). Together, these observations suggest that fast pDMS value coding is aligned with pre-existing short-latency sensory responses present before learning, whereas anterior ventral value coding reflects a longer-latency CS+ preferring response recruited after learning.

Given that a subset of pDMS sites encoded both cue value and cue location, we next asked whether learned value signals were expressed uniformly across cue positions or interacted with pre-existing location selectivity. In principle, value and location could be encoded independently, such that CS+ and CS- differences remain similar across cue positions, or interactively, such that value encoding depends on stimulus location (Supplementary Fig. 5A). To explore these possibilities, we examined a subset of pDMS sites that showed both value coding and location selectivity. We observed examples consistent with value-location interactions in both pathways (Supplementary Fig. 5B,C). Among dual-encoding sites, a subset showed a significant value × location interaction, including 3/12 dSPN sites and 4/11 iSPN sites (linear mixed-effects model as a function of location, value and the interaction, with mouse included as random intercept, p < 0.05; Methods). Notably, the interaction patterns were consistent with the pathway-specific value preference observed in pDMS: interacting dSPN sites preferentially encoded contralateral CS+ cues, whereas interacting iSPN sites preferentially encoded contralateral CS- cues. Although this analysis was limited by the small number of dual-encoding sites, these observations raise the possibility that learned value signals in pDMS can interact with pre-existing location selectivity rather than being added uniformly across cue locations.

Taken together, these results reveal two distinct forms of striatal cue-value representations: shorter-latency, pathway-opponent encoding in pDMS and longer-latency CS+ preferring encoding shared by dSPNs and iSPNs in anterior ventral striatum.

### Temporal reward proximity is encoded by dSPN-biased ramping in posterior ventrolateral striatum

In addition to short-latency cue-evoked Ca^2+^ responses, CS+ trials elicited a slower monotonic response component that increased across the delay period and peaked near the time of reward delivery (Fig. 4A). This ramping was largely absent during CS- trials. To quantify monotonic ramping, we fit each recording site with an orthogonal time-basis model designed to separate linear ramping components from nonramping cue-evoked response dynamics (Methods; Supplementary Fig. 7A). This approach distinguished sites with sustained monotonic increases from sites whose trial-averaged activity appeared ramp-like but was better explained by higher-order components, such as temporally variable transient activity (Supplementary Fig. 7B,C). Ramping signals were strikingly pathway specific, with significantly more ramping sites present in dSPNs than iSPNs (75/364, 20.6% dSPN sites and 13/329, 4.0% iSPN sites; linear mixed model of ramping significance as a function of cell type with random intercept for mouse, p = 1.83 × 10^-5^). Moreover, dSPN ramping responses were concentrated in posterior ventrolateral striatum (pVLS; Fig. 4B). Thus, reward-predictive cues recruited a spatially localized, dSPN-biased ramping signal during the delay period.

**Figure 4.**
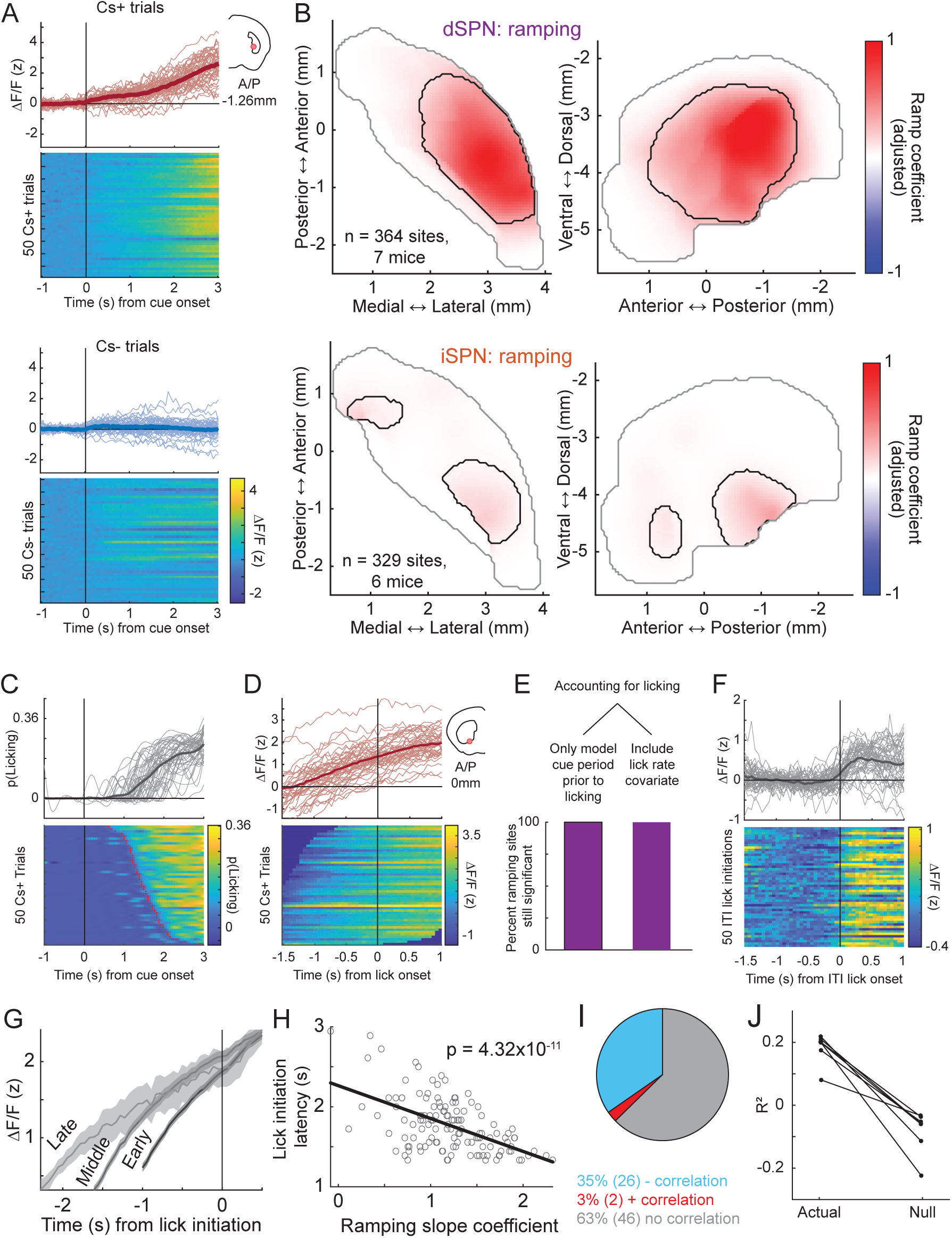
Temporal reward proximity is represented by dSPN-selective ramping in posterior ventrolateral striatum. (A) Example dSPN ramping site. Top, trial-by-trial activity and mean activity for 50 CS+ trials and 50 CS- trials. Thin lines indicate individual trials, and thick lines indicate trial averages. Coronal schematic indicates site location. Bottom, corresponding trial-by-trial activity heat maps. (B) Smoothed horizontal and sagittal maps of ramping strength in dSPN and iSPN datasets. Voxel color indicates the linear ramp coefficient after spatial smoothing, with red indicating positive ramping. Black contours are significant ramping regions. dSPN sites showed a higher fraction of ramping sites than iSPN sites. Generalized linear mixed model of ramping significance as a function of cell type with mouse as a random effect; p = 1.83 × 10 ^-5^. (C) Trial-by-trial prelicking behavior aligned to cue onset. Top, licking probability on individual trials and mean licking probability. Thin lines indicate individual trials, and thick line indicates the trial average. Bottom, corresponding trial-by-trial heat map of licking activity. (D) Trial-by-trial calcium activity from an example dSPN site aligned to first prelick initiation. Top, individual trials and mean activity. Thin lines indicate individual trials, and thick line indicates the trial average. Bottom, corresponding trial-by-trial heat map of calcium activity. Coronal schematic indicates site location. (E) Percent of dSPN ramping sites that remained significantly ramping after controlling for licking in two ways: restricting the model to the cue period before lick initiation or including lick rate as a continuous covariate. (F) Same example dSPN site as in (D), aligned to spontaneous lick-bout onsets during the inter-trial interval. Top, individual trials and mean activity. Thin lines indicate individual trials, and thick line indicates the trial average. Bottom, corresponding trial-by-trial heat map of calcium activity aligned to ITI lick initiations. (G) Mean activity from the example dSPN site in (D), grouped by lick initiation latency. Trials were grouped into early, middle, and late lick-initiation tertiles. Shaded regions indicate SEM. (H) Relationship between ramp slope and lick initiation latency on each trial for the example site in (D). Each dot represents one trial. Steeper ramping was associated with earlier prelick initiation; p = 4.32 × 10^-11^. (I) Fraction of dSPN ramping sites showing a negative correlation, positive correlation, or no significant correlation between ramp slope and lick initiation latency across trials. Counts indicate numbers of recording sites. (J) Prediction of lick initiation latency from ramp slope for each mouse compared to predictions obtained from a null distribution. Each line represents one mouse. Statistical significance was assessed using a one-sided empirical permutation test, in which trial labels were shuffled 500 times and the p value was computed as the fraction of shuffled R^2^ values greater than or equal to the observed mean R^2^. For (B), learned-stage datasets include dSPNs, n = 364 sites from 7 mice, and iSPNs, n = 329 sites from 6 mice. For (E)-(J), analyses were restricted to dSPN ramping sites.

Because CS+ trials also evoked anticipatory licking, we next asked whether dSPN ramping could instead reflect lick execution or lick preparation. Several observations argued against this interpretation. First, ramping was consistent across trials and began before the first lick, indicating that it was not simply a signal aligned to lick execution (Fig. 4C,D). Second, accounting for licking in the model did not reduce the ramping signal. Monotonic ramping coefficients remained significant when lick rate was included as a continuous covariate in the orthogonal time-basis model and when the analysis was restricted to the cue period before each trial’s first lick (Methods; Fig. 4E; Supplementary Fig. 8A,B). Third, comparable ramping activity did not precede spontaneous lick-bout onsets during the inter-trial interval, indicating that ramps do not reflect a general lick-preparatory or pre-motor signal (Fig. 4F). Of the 75 dSPN sites with significant delay-period ramping, only 8 sites (10.7%) exhibited monotonic ramping before ITI lick initiation. Though dSPN ramping was dissociable from lick execution dynamics, signals linked to lick execution were present within the same pVLS region, consistent with prior anatomical and functional studies^2,10,47,51^. Lick-locked Ca^2+^ signaling was present in both dSPN and iSPN recordings within this region (51/65 dSPN sites and 57/82 iSPN sites), unlike the strongly dSPN-biased ramping signal (Supplementary Fig. 9A,B). Together, these analyses indicate that dSPN ramping is not explained by lick execution or nonspecific motor preparation, even though lick-related encoding is co-expressed within the same pVLS region across both pathways.

Although ramping did not directly reflect licking per se, its trial-by-trial dynamics could still relate to when anticipatory licking was initiated. If ramping reflects an internal estimate of temporal reward proximity, then steeper ramps might bring the animal more rapidly toward a behavioral threshold for anticipatory licking. Consistent with this possibility, trials with earlier lick initiation had steeper pre-lick increases in dSPN activity (Fig. 4G). Across individual trials, ramp slope was negatively correlated with lick initiation latency, such that steeper ramping predicted earlier anticipatory licking (Fig. 4G,H; linear regression, p = 4.32 × 10^-11^). Across the population, many dSPN ramping sites showed this negative trial-by-trial relationship (Fig. 4I), and at the mouse level, ramp slopes predicted lick initiation latency better than expected from a null distribution (Methods; Fig. 4J; empirical p = 0.002 in all mice, computed as the proportion of shuffled R^2^ values greater than or equal to the observed R^2^). Thus, while dSPN ramping was not reducible to lick execution or motor preparation, its slope carried information about the timing of anticipatory licking.

### Glutamatergic input dynamics distinguish region-specific inherited and locally transformed striatal signals

The region- and cell-type-specific signals observed in SPN calcium recordings could arise through at least two mechanisms. In one model, SPN calcium dynamics inherit task-related selectivity from afferent inputs that already encode cue location, cue value, or reward timing. Alternatively, glutamatergic inputs may provide sensory or task-related signals that are transformed locally within striatal circuits into pathway-specific SPN dynamics (Fig. 5A). To distinguish between these possibilities, we measured striatum-wide glutamate release dynamics using flex-iGluSnFR targeted to dSPNs or iSPNs in separate D1-Cre and A2A-Cre mice (dSPN-targeted recordings: n = 3 mice; iSPN-targeted recordings: n = 4 mice; Fig. 5B; Supplementary Fig. 10). This approach revealed widespread cue-evoked glutamate responses across the striatum (dSPN-targeted sites: 99/215; iSPN-targeted sites: 187/242; Fig. 5C), which were largely absent in quasi-simultaneous isosbestic control recordings (dSPN-targeted sites: 8/215; iSPN-targeted sites: 7/242).

**Figure 5.**
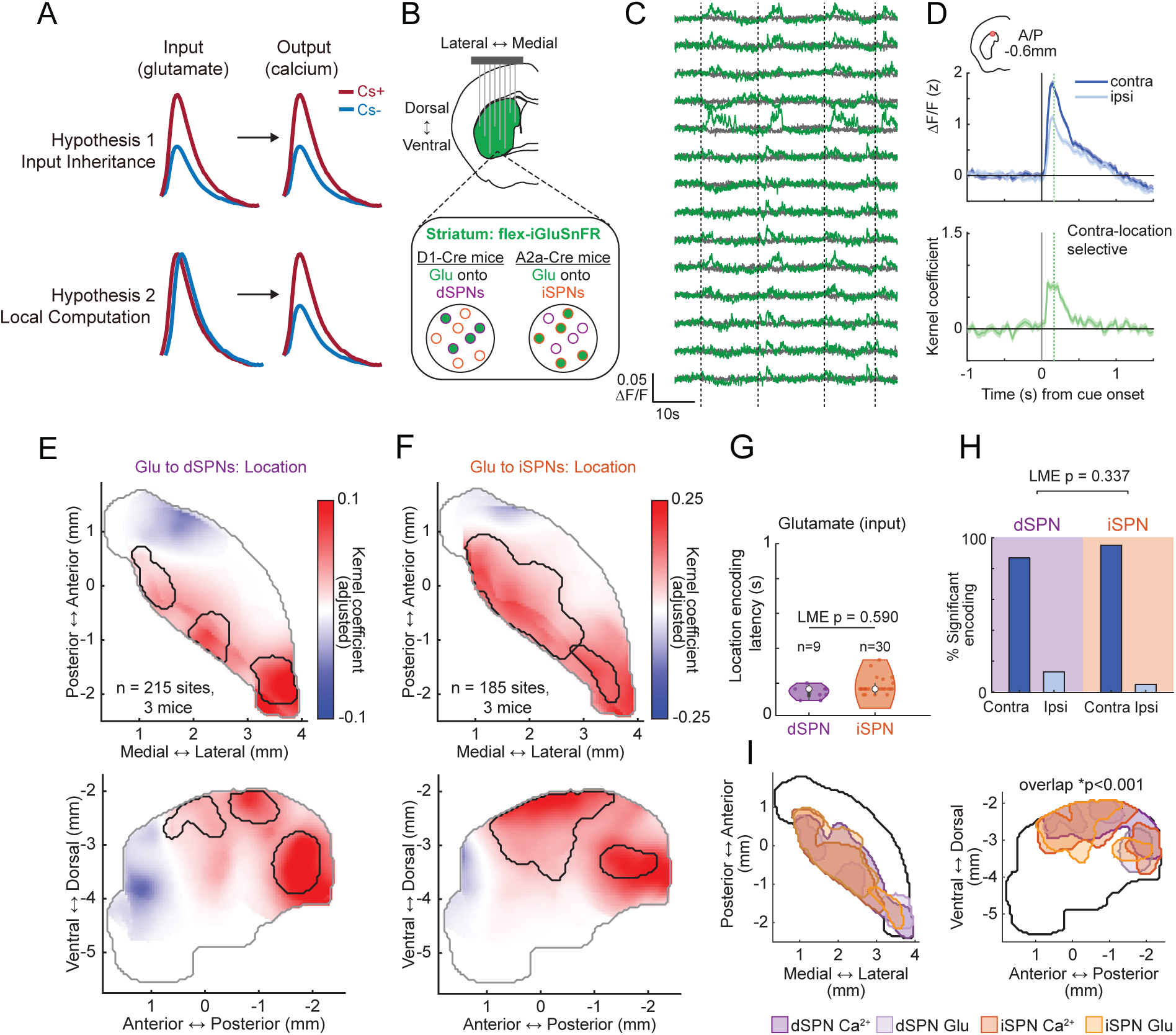
Cue-location encoding is present in short-latency pDMS glutamatergic input. (A) Conceptual models distinguishing inherited input selectivity from local computation. Under the input-inheritance model, SPN calcium selectivity matches glutamatergic input selectivity. Under the local-computation model, SPN calcium selectivity can diverge from glutamatergic input selectivity. (B) Schematic of the implantation and glutamate-sensor targeting strategy used to record glutamatergic input to dSPNs and iSPNs. flex-iGluSnFR expression was restricted to dSPNs or iSPNs in separate D1-Cre and A2A-Cre cohorts. (C) Representative simultaneously recorded iSPN-targeted glutamate fluorescence signals across sites from one session. Each row corresponds to one recording site. Green traces show glutamate-dependent 470 nm flex-iGluSnFR signals and gray traces show 405 nm isosbestic control traces. Dashed vertical lines indicate cue presentations. (D) Example iSPN glutamate-recording site with significant cue-location selectivity. Top, mean ΔF/F responses to contralateral and ipsilateral cue presentations. Shaded regions indicate SEM. Bottom, model kernels for the same site. Green vertical line indicates the time at which the location-selective kernel reached 80% of peak amplitude. (E) Smoothed horizontal and sagittal maps of glutamatergic cue-location encoding in dSPN-targeted recordings. Voxel color indicates the location-kernel coefficient amplitude after spatial smoothing, with red indicating contralateral preference and blue indicating ipsilateral preference. Black contours mark significant group-level location-encoding regions. (F) Same as (E), for iSPN-targeted glutamate recordings. (G) Distribution of location-encoding latencies for glutamatergic input to dSPNs and iSPNs within the significant location encoding contour. Dots indicate individual recording sites. Location-encoding latency did not differ between cell types. Linear mixed model with latency as a function of cell type and mouse as a random effect; p = 0.590. (H) Cue-location preference among location-selective glutamate-recording sites. Bars indicate the percentage of significant location-encoding sites preferring contralateral or ipsilateral cue locations. Contra/ipsi preference did not differ between dSPN- and iSPN-targeted recordings. Generalized linear mixed-effects model with significance status predicted by cell type and encoding preference and mouse as a random effect; p = 0.337. (I) Overlay of significant location-encoding regions for dSPN and iSPN calcium and glutamate recordings. The four regions significantly overlapped (p < 0.001). For glutamate recordings, dSPN-targeted datasets include n = 215 sites from 3 mice and iSPN-targeted datasets include n = 185 sites from 3 mice for location analyses unless otherwise indicated. Spatial overlap was assessed using the spatial overlap test described in Methods. Asterisks indicate significant time points in model kernels: *p < 0.05; ns, not significant.

We first asked whether the short-latency pDMS cue-location signal observed in SPN calcium activity was also present in glutamatergic input. Using the same regression approach applied to calcium recordings, we isolated location-selective components of cue-evoked glutamate release (Fig. 5D). Cue-location selectivity was detected in a subset of glutamate-recording sites in both dSPN and iSPN targeted mice (dSPN-targeted sites: 15/215; iSPN-targeted sites: 38/185; Fig. 5D-F). Most location-selective glutamate sites favored contralateral cue locations (dSPN-targeted sites: 13/15; iSPN-targeted sites: 36/38), and this preference did not differ between dSPN- and iSPN-targeted recordings (linear mixed-effects model as a function of location with random intercept for mouse, p = 0.337; Fig. 5H). Location-encoding latencies were similarly short in both targeted populations (linear mixed-effects model as a function of cell type with random intercept for mouse, p = 0.590; Fig. 5G), and significant glutamate location-encoding regions were concentrated in pDMS, closely overlapping the dSPN and iSPN calcium location-encoding regions (spatial overlap test, p < 0.001; Fig. 5E,F,I). Thus, the rapid pDMS cue-location signal was present in glutamatergic input and shared across SPN-targeted recordings, consistent with inheritance of a sensory spatial signal by both pathways.

We next asked whether glutamatergic input also contained the pathway-divergent value signals observed in pDMS calcium activity. Significant value encoding was present in many glutamate-recording sites in both dSPN and iSPN targeted mice (dSPN-targeted sites: 55/215; iSPN-targeted sites: 121/242; Fig. 6A-C), indicating that cue-value information was present in striatal afferent input. However, unlike calcium signals, in which iSPN sites frequently showed CS- preference in pDMS, glutamatergic value encoding was almost uniformly CS+ preferring in both dSPN and iSPN targeted recordings (54/55 dSPN-targeted sites; 112/121 iSPN-targeted sites; Fig. 6A-D). CS+ preferring glutamate signals were strongest in anterior ventral striatum and extended into anterior dorsal striatum, whereas value encoding was weak or absent in the pDMS region where calcium signals showed pathway-opponent value coding (Fig. 6B,C). Within the divergent pDMS value-coding region, glutamate value-kernel amplitudes did not differ between dSPN- and iSPN-targeted recordings, in contrast to the strong pathway divergence observed in calcium dynamics (linear mixed-effects model as a function of cell type and sensor with random intercept for mouse; glutamate, p = 0.326; calcium, p = 8.78 × 10^-7^; Fig. 6D). Thus, pathway-opponent pDMS value coding was not directly inherited from similarly opponent glutamatergic input, but instead reflected a divergence between afferent input and SPN calcium dynamics.

**Figure 6.**
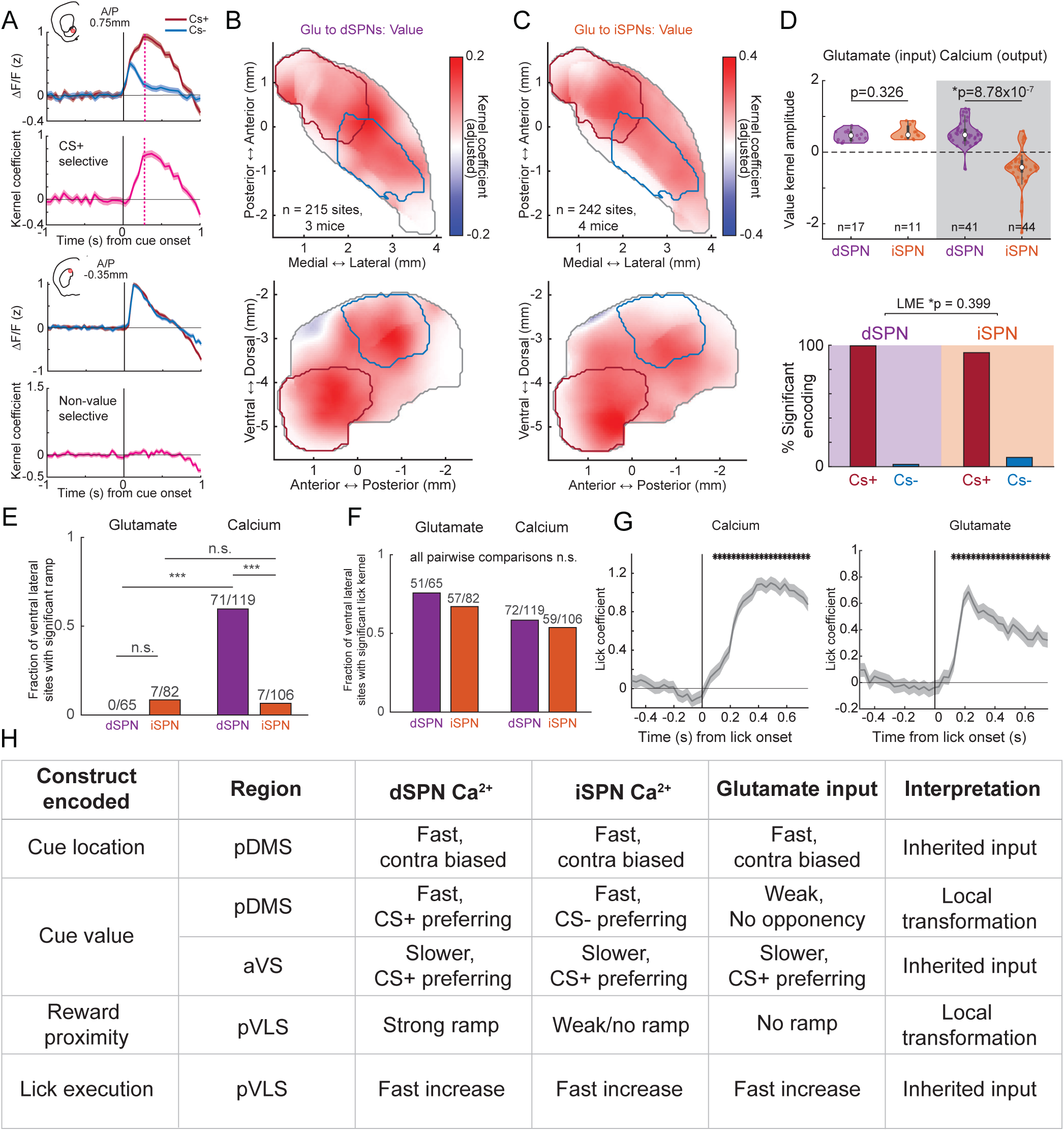
SPN value and reward-proximity signals diverge from glutamatergic input. (A) Example iSPN glutamate-recording sites. Top example, mean .LF/F responses to CS+ and CS- cues and corresponding model kernels for an anterior ventral site with value selectivity (CS+ preference). Bottom example, same as top for a pDMS site with no value selectivity. Magenta traces show value-selective kernels. Magenta vertical lines indicate the time at which the value-selective kernel reached 80% of peak amplitude. Shaded regions indicate SEM. (B) Smoothed horizontal and sagittal maps of glutamatergic cue-value encoding in dSPN-targeted recordings. Voxel color indicates the value-kernel coefficient amplitude after spatial smoothing, with red indicating CS+ preference and blue indicating CS- preference. Red and blue contours denote the convergent and divergent calcium value-encoding regions, respectively. (C) Same as (B), for iSPN-targeted glutamate recordings. (D) Top, value-kernel coefficients within the divergent calcium value-encoding region. Violin plots show value-kernel amplitudes for glutamatergic input and calcium signaling in dSPN and iSPN datasets. Dots indicate individual recording sites. Glutamatergic value-kernel amplitudes did not differ between dSPN- and iSPN-targeted recordings, whereas calcium value-kernel amplitudes differed between dSPNs and iSPNs. Generalized linear mixed model with kernel amplitude as a function of recording type and cell type and mouse as a random effect; glutamate dSPN vs iSPN, p = 0.326; calcium dSPN vs iSPN, p = 8.78 × 10^-7^. Bottom, cue-value preference among value-selective glutamate-recording sites. CS+/CS- preference did not differ between dSPN- and iSPN-targeted recordings. Linear mixed-effects model with significance status predicted by cell type and encoding preference and mouse as a random effect; p = 0.399. (E) Fraction of sites in the ventrolateral ramping region with significant monotonic ramping. Ramping was largely absent from glutamate recordings but enriched in dSPN calcium recordings. Numbers above bars indicate significant ramping sites over total sites. Generalized linear mixed-effects model with significance status predicted by recording type and cell type and mouse as a random effect. (F) Fraction of sites within the ventrolateral ramping region with significant lick-related kernels across dSPN-targeted and iSPN-targeted glutamate recordings and dSPN and iSPN calcium recordings. Numbers above bars indicate significant lick-encoding sites over total sites. The fraction of lick-encoding sites did not differ across recording type or cell type. Generalized linear mixed-effects model with significance status predicted by recording type and cell type and mouse as a random effect; all pairwise comparisons ns. (G) Lick-initiation kernels estimated from one example iSPN calcium recording and one example iSPN glutamate recording from posterior ventrolateral striatum. Shaded region indicates SEM. Asterisks mark time points with significant lick-related encoding. (H) Summary of cue-location, cue-value, and reward-proximity encoding across dSPN calcium, iSPN calcium, and glutamatergic input recordings. The table summarizes whether each signal was consistent with inherited glutamatergic input or local transformation from input to SPN calcium dynamics. For glutamate recordings, dSPN-targeted datasets include n = 215 sites from 3 mice and iSPN-targeted datasets include n = 242 sites from 4 mice unless otherwise indicated. Calcium datasets are as described in Figures 1-4. Asterisks indicate significant time points or pairwise comparisons: *p < 0.05, ***p < 0.001; ns, not significant.

Finally, we asked whether the dSPN-biased ramping response in pVLS was present in glutamatergic input. Ramping was largely absent from glutamate release dynamics within the ventrolateral ramping region, with no ramping sites detected in dSPN-targeted glutamate recordings and only a small number detected in iSPN-targeted glutamate recordings (dSPN-targeted sites: 0/65; iSPN-targeted sites: 7/82; Fig. 6E). This absence was not due to a general inability to detect behavior-related glutamatergic signals in this region. Lick-related glutamate signals were readily detected in posterior ventral striatum, including within the pVLS region containing dSPN calcium ramping (Fig. 6F,G; Supplementary Fig. 9C,D), similar to lick-related calcium signals observed in the same territory (Supplementary Fig. 9A,B). Thus, glutamatergic inputs to pVLS carried lick-related activity but did not transmit the dSPN-biased monotonic ramping observed in calcium dynamics.

Together, these input-output comparisons revealed that striatal task signals differed in their relationship to glutamatergic input (Fig. 6H). Cue-location encoding in pDMS closely matched glutamatergic input and was shared across SPN pathways. In contrast, pathway-opponent pDMS value coding and dSPN-biased pVLS ramping were present in SPN calcium dynamics but not in measured glutamatergic input. These results support a model in which some striatal signals are inherited from afferent input, whereas others emerge through region and pathway-specific transformations from input to SPN calcium dynamics.

## Discussion

Using large-scale recordings of population calcium signals within distinct SPN subtypes, we revealed a spatiotemporal organization of perceptual and motivational task features across the striatum. During Pavlovian visual conditioning, cue onset, cue location, learned cue value, reward proximity, and lick-related behavior were represented in distinct anatomical territories and time windows. Some of these representations were shared across dSPNs and iSPNs, whereas others were pathway-divergent or pathway-opponent, indicating that striatal task representations are organized not only by region and time, but also by SPN pathway. By comparing these calcium dynamics with targeted measurements of glutamatergic input, we found that rapid cue-location signals and lick-related signals were also present in glutamatergic input dynamics, consistent with inherited sensory and behavioral representations. In contrast, pathway-opponent pDMS value signals and dSPN-biased pVLS ramping diverged from measured glutamatergic input, suggesting that learning-dependent, cell-type-specific dynamics can emerge through local striatal transformations. Together, these findings identify a regionally organized input-output architecture in which striatal circuits inherit some task-related signals while transforming others into pathway-specific representations of learned value and reward proximity.

A central implication of these results is that learning-related striatal representations may be shaped by the pre-existing input structure present in each region. In pDMS, short-latency visual responses and cue-location selectivity were already present before conditioning and were expressed similarly in dSPNs, iSPNs, and glutamatergic inputs. This inherited sensory structure may provide a scaffold that constrains where and how learning-dependent plasticity is expressed. After conditioning, this same posterior dorsomedial territory contained pathway-divergent value signals, with dSPNs preferentially responding to the reward-predictive CS+ cue and iSPNs preferentially responding to the non-rewarded CS- cue. In contrast, anterior striatal regions, particularly the ventromedial striatum, exhibited little cue-evoked activity before learning but acquired longer-latency CS+ preferring responses in both dSPNs and iSPNs after learning. This uniform CS+ preference was also present in glutamatergic inputs, suggesting that value-related activity in this region may be inherited, at least in part, from learned responses acquired in distributed cortical and subcortical afferents, including prefrontal cortex, amygdala, thalamus, and hypothalamus. This does not imply that local plasticity is absent in ventral striatum or that dSPNs and iSPNs are functionally equivalent in this region. However, it suggests that any local pathway-specific transformations may be layered onto already value-selective input dynamics, rather than built directly from a pre-existing sensory scaffold. More generally, these results support a model in which afferent inputs define the variables available to a given striatal region, whereas local plasticity and circuit mechanisms determine how those variables are converted into SPN dynamics within distinct cell types.

What is the mechanism for the divergent pDMS value signals? One possibility is that dopamine-dependent plasticity modifies pre-existing visual inputs onto dSPNs and iSPNs according to distinct pathway-specific rules. Because dSPNs and iSPNs express different dopamine receptor classes and are subject to distinct forms of dopaminergic modulation, reward-predictive teaching signals could strengthen cue-evoked responses in one pathway while suppressing, reshaping, or selectively gating responses in the other^6,7,9,52^. However, pathway divergence could also emerge through local filtering of otherwise similar input, including differences in dendritic integration, feedforward inhibition, or cholinergic modulation^35,36,53^. Our data cannot distinguish among these mechanisms, but they argue against the simplest inheritance model because glutamatergic input to pDMS did not show the pathway-opponent CS+ and CS- preferences observed in SPN calcium signals. Thus, the divergent pDMS representations likely reflect local processing within striatal circuitry rather than direct transmission of an already divergent upstream value signal. Although our findings support a broadly similar topography of glutamatergic input encoding across dSPN and iSPN populations, it is possible that pathway-specific input dynamics differ in other task conditions, in specific subcellular compartments, or in more subtle ways than our cross-mouse population measurements could detect. Indeed, tracing and anatomical studies indicate that dSPNs and iSPNs receive broadly overlapping but not identical glutamatergic inputs ^33,54^.

The functional role of these pDMS signals is also unresolved, but their combination of spatial and value selectivity suggests that they may contribute to assigning motivational significance to cue locations. One possibility is that CS+ preferring dSPN activity promotes orienting or attentional priority toward valued cues, whereas CS- preferring iSPN activity promotes disengagement from cues that do not predict reward^38,55–57^. This interpretation is consistent with the localization of these signals to a region containing rapid visual and location-selective responses, and with the observation that value and location coding can interact in individual pDMS sites. Although speculative, this hypothesis provides a plausible functional framework for why pathway divergence would emerge from a pre-existing spatial sensory scaffold. Testing this possibility will require targeted perturbations of pDMS dSPNs and iSPNs during CS+ and CS-presentations to determine whether these signals causally influence cue engagement, orienting, or suppression of responses to non-rewarded cues.

In contrast to rapid pDMS value signals, reward-predictive cues also elicited a slower ramping signal localized to posterior ventrolateral striatum. This ramping signal, which reflected temporal reward proximity, was present almost exclusively in dSPNs and was largely absent during CS-trials. Although ramping was expressed in a region containing dSPN and iSPN signals linked to licking, it was not explained by measured lick-related activity or motor preparation per se as it did not precede spontaneous licking outside the task and persisted after accounting for lick execution. Nevertheless, we cannot exclude the possibility that unmeasured covert orofacial movements, postural adjustments, or other preparatory motor variables covary with reward proximity and contribute to the ramping signal. Importantly, however, the slope of the dSPN ramp predicted the subsequent timing of anticipatory lick initiation, suggesting that the signal is linked to the evolving probability or readiness for anticipatory responding as reward delivery approaches. In this view, the ramp does not deterministically trigger licking, but increasing dSPN activity may bring the animal closer to a behavioral activation threshold, with steeper ramps leading to earlier anticipatory licking. This interpretation is conceptually similar to accumulator or drift-to-threshold models widely used to explain the timing of decisions and actions ^58–61^, and is consistent with anatomical and behavioral evidence linking ventrolateral striatal circuits to licking and orofacial control ^10,47,62^. Directly inhibiting or imposing ramp-like activity in pVLS dSPNs during the delay period could test whether this signal causally controls the timing or probability of anticipatory licking.

The scaffold for this pVLS ramping signal is less clear than for pDMS value coding. Unlike pDMS, where cue-location selectivity and early visual responses were evident before learning and matched by glutamatergic input, monotonic ramping was not observed in glutamatergic input to either dSPNs or iSPNs. This absence is unlikely to reflect a general inability to detect behaviorally relevant glutamate signals, because lick-related glutamate dynamics were readily detected in the same posterior ventral territory. Nevertheless, differences in sensor kinetics, signal integration, or spatial averaging could make some forms of temporally structured glutamatergic input difficult to detect, particularly if they are sparse or distributed across sequentially active inputs. One possibility is that pVLS contains or receives an interval-specific representation of elapsed time from cue onset, encoded within temporally structured SPN ensembles, which is transformed locally into a dSPN-biased ramping signal during learning. In this model, dSPN inputs or ensembles active near the expected time of reward could be preferentially strengthened, perhaps through reward-evoked dopamine, resulting in a population-level ramp after learning. Such temporally structured ensemble activity could be obscured in bulk measurements, but sequential timing activity has been observed in striatal SPN populations^63–65^. A second possibility is that ramping dopamine signals are transmitted to pVLS and preferentially increase dSPN excitability during the delay period. Dopamine ramps have been observed in striatum, particularly in ventral territories, during approach to reward and action-sequence execution^66–69^. Moreover, slowly evolving dopamine activity before self-timed movements can predict and bias movement timing^70^. However, whether such dopamine ramps are present in pVLS during this task, and whether they are sufficient to generate the observed dSPN-biased calcium ramping, remains unknown. Distinguishing between these possibilities will require direct measurements and manipulations of dopamine, glutamatergic afferents, and SPN activity during learning.

A limitation of these experiments is that fiber photometry reports population calcium dynamics and cannot resolve how signals are distributed across individual neurons, dendrites, axons, or local neuropil. This is particularly relevant in the striatum, where bulk photometry signals may differ from individual somatic activity and can include nonsomatic or neuropil-associated calcium dynamics that complicate interpretation of population signals^71^. However, several observations argue against a simple interpretation in which the key pathway-specific signals measured here primarily reflect excitatory input dynamics. First, targeted glutamate recordings showed that SPN calcium signals did not always mirror local glutamatergic input. Rapid cue-location encoding in pDMS was closely matched between glutamate and calcium signals, whereas pathway-opponent pDMS value signals and dSPN-biased pVLS ramping were absent from corresponding glutamatergic input dynamics. In addition, concurrent photometry from striatal SPN populations and their downstream axonal projections has shown that striatal calcium transients are strongly reflected in axonal activity, supporting the interpretation that pathway-specific SPN population calcium dynamics can capture output-relevant activity^72^. Thus, although cellular-resolution imaging and electrophysiology will be needed to determine how these population signals arise from heterogeneous SPN ensembles, their pathway specificity, anatomical organization, and divergence from glutamatergic input indicate that the signals identified here reflect structured SPN population dynamics that cannot be explained by local excitatory input alone.

In conclusion, our findings support a model in which learned striatal population signals emerge when local circuit mechanisms transform regionally organized sensory, temporal, or motor scaffolds into pathway-specific activity patterns. In this framework, glutamatergic input topography determines which variables are available for transformation in each striatal territory, and local plasticity and neuromodulation determine whether those variables become expressed as dSPN/iSPN imbalances. The behavioral function of each transformed signal may then depend on the output channels and action programs associated with that territory. Temporal reward-proximity signals in pVLS may support the appropriate timing of licking through ventrolateral circuits linked to orofacial control, whereas spatial value signals in pDMS may support orienting, engagement, or disengagement with valued and non-valued cues through circuits involved in spatial attention and locomotor orienting. Thus, learning may not impose a single global value representation across the striatum, but instead selectively modifies local input-defined scaffolds to generate cell-type-specific output signals deployed at the appropriate time, in the appropriate context, and toward the appropriate stimulus.

## Methods

### Animals

Adult male and female Drd1-Cre transgenic mice (MMRRC, strain #037156-JAX; n = 3 males and 5 females; postnatal age 8-25 weeks; body weight 24-30 g) and Adora2a-Cre mice (MMRRC, strain #4361654; n = 4 males and 2 females; postnatal age 8-25 weeks; body weight 24-30 g) were used for cell-type-specific expression of the calcium indicator^73^. For expression of the glutamate indicator, a separate cohort of adult Drd1-Cre mice (MMRRC, strain #037156-JAX; n = 1 male and 2 females) and Adora2a-Cre mice (MMRRC, strain #4361654; n = 2 males and 2 females) was used.

Mice were housed on a 12 h light-dark cycle, with lights on from 9 PM to 9 AM, and all experiments were performed during the dark cycle. Before surgery, mice were group-housed with 2-5 animals per cage. After implantation, mice were housed individually. During behavioral training and imaging experiments, mice were water-restricted to 1 mL per day, adjusted as needed to maintain 80-90% of their initial body weight. Food was available ad libitum. All procedures were approved by the Boston University Institutional Animal Care and Use Committee.

### Multi-fiber array fabrication and calibration

Multi-fiber arrays were fabricated in-house to enable large-scale fiber photometry recordings across deep brain volumes. The detailed fabrication procedure has been described previously^51^. Briefly, optical fibers (Fiber Optics Technology) with outer diameters of either 37 µm (34 µm core, 3 µm cladding) or 50 µm (46 µm core, 4 µm cladding) and a numerical aperture of 0.66 were used. Fibers were inserted under a microscope into 55-60 µm diameter holes in a custom 3D-printed grid (3 mm width × 5 mm length; Boston MicroFabrication). Fiber lengths were measured under a dissection microscope to target specific depths below the grid and were then secured using UV-curable adhesive (Norland Optical Adhesive 61). The distal, implanted ends of the fibers were arranged according to the target striatal sampling pattern, whereas the proximal ends were bundled within an approximately 1 cm section of polyimide tubing (0.8-1.3 mm diameter; MicroLumen). The proximal fiber bundle was cut with a fresh razor blade and polished using fine-grained polishing paper (Thorlabs) to produce a smooth and uniform imaging surface. Each array contained 60-110 fibers, with a minimum spacing of 220 µm along the medial-lateral axis and 250 µm along the anterior-posterior axis. These separations were chosen to maximize coverage of the striatal volume while preventing overlap between the optical collection fields of individual fibers^51^. A larger-diameter post was attached to one side of the plastic grid to facilitate handling during implantation. Before implantation, each array was calibrated to map the implanted fiber positions to their corresponding locations on the proximal imaging surface. For calibration, the implanted fiber tips were illuminated while transmitted light was imaged at the bundle surface. By sequentially illuminating different rows and columns of fibers, each fiber observed at the imaging surface was matched to its row and column position on the grid and to its corresponding distal fiber tip.

### Surgical viral injection & array implantation

Genetically encoded fluorescent indicators for optical recording were expressed by injection of adeno-associated viruses (AAVs). For calcium recordings, a Cre-dependent viral vector encoding the green fluorescent calcium indicator jGCaMP7f (AAV1.hSyn.FLEX.jGCaMP7f.WPRE, 3.60 × 10^12^ to 7.00 × 10^12^ GC/mL; Addgene) was used^74^. In Drd1-Cre and Adora2a-Cre mice, this approach enabled selective expression of the calcium indicator in direct-pathway spiny projection neurons (dSPNs) and indirect-pathway spiny projection neurons (iSPNs), respectively. For glutamate recordings, a Cre-dependent viral vector encoding the green fluorescent glutamate sensor iGluSnFR (AAV1.hSyn.FLEX.SF-iGluSnFR.A184S, 2.50 × 10^12^ to 4.00 × 10^12^ GC/mL; Addgene) was used^75^. In Drd1-Cre and Adora2a-Cre mice, this approach enabled cell-type-specific expression of membrane-localized iGluSnFR in dSPNs or iSPNs, allowing detection of local glutamate transients near targeted SPN membranes.

Mice were anesthetized with isoflurane (1-3%) and placed in a stereotaxic frame. A large craniotomy was made using a surgical drill (Midwest Tradition 790044, Avtec Dental RMWT) to expose the cortical surface above the striatum. Virus was injected stereotaxically into the striatum through a pulled glass pipette with a 30-50 µm tip diameter. In total, 30-40 injection sites were targeted across the striatum to maximize overlap between viral expression and the planned fiber positions. At each site, 200-400 nL of virus was delivered at approximately 100 nL/min. Following viral injection, the multi-fiber array was mounted in the stereotaxic apparatus for implantation. The longest fiber in the array served as a calibration fiber and was aligned to bregma before being moved to its designated stereotaxic coordinate. The array was then slowly lowered into the brain to the target depth. After implantation, a thin layer of Kwik-Sil (WPI) was applied around the craniotomy to seal the exposed edges. Tissue adhesive (3M VetBond) was then used to attach the implant edges to the Kwik-Sil layer to prevent cerebrospinal fluid leakage. Metabond (Parkell) was subsequently applied to secure the plastic grid of the fiber array to the skull. Once the initial layer of Metabond had cured, a metal headplate and ring (Atlas Tool and Die Works) were attached to the skull using additional Metabond, and the exposed surface was coated with blackened Metabond mixed with carbon powder (Sigma) to reduce light contamination. Finally, a cylindrical plastic protective cap, trimmed to extend just above the end of the fiber bundle, was secured around the array and lined internally with blackened Metabond.

Behavioral habituation began at least one week after surgery, and neural data acquisition began 3-5 weeks after viral injection and array implantation to permit expression of the fluorescent sensors.

### Head-fixed behavior apparatus

Mice were head-fixed with their limbs resting on a hollow 8-inch diameter Styrofoam ball that floated on a cushion of pressurized air within a 3D-printed plastic cradle. The cradle contained small air vents that supported smooth ball rotation during locomotion. Ball movement was measured along three rotational axes, pitch, yaw, and roll, using two optical mouse sensors with the outer shell removed (Logitech G203). The sensors were positioned at the equator of the ball and oriented orthogonally to capture different rotational components. One sensor recorded pitch and yaw rotation, whereas the second sensor measured roll rotation. Both sensors were configured at 400 dpi with a polling rate of 1 kHz. Sensor outputs were processed by a Raspberry Pi 3B+, which converted rotational signals into analog voltage signals at 100 Hz. A running velocity of 3.5 m/s corresponded to an output voltage of 3.3 V. These signals were relayed to a data acquisition board (National Instruments PCIe-6343). Running direction was simultaneously encoded as a digital binary signal from the Raspberry Pi and transmitted through the same data acquisition system.

Water rewards were delivered through a 60 mL syringe connected by plastic tubing to a digitally gated solenoid valve (Neptune Research, #161T012). The valve controlled the flow of water to a metal spout mounted in front of the animal. Licking behavior was detected using a capacitive touch sensor attached to the spout.

Behavioral signals, including velocity, licking, and task-related TTLs, were acquired at 2 kHz using a custom MATLAB program and the National Instruments data acquisition board. The same system controlled task outputs, including reward delivery, LED triggering, and image acquisition. For synchronization of behavioral and imaging data, TTL pulses were sent from the cameras to the acquisition board 500 µs after the beginning of readout for each frame (Hamamatsu HCLive VSYNC). Behavioral data were downsampled to match the sampling rate of the neural data by block averaging within each imaging frame. Binary behavioral variables were additionally downsampled by rounding or summing, depending on the analysis.

### Three-location Pavlovian task

Visual cues were generated using the virtual reality MATLAB engine ViRMEn^76^ and displayed on an array of five 12 × 21 inch monitors arranged in a semicircle approximately 13 inches in front of the mice. Two cue identities were used, consisting of vertically or horizontally oriented black-and-white stripes extending approximately 10 inches in width. One cue was assigned as the reward-predictive cue (CS+) and the other as the non-rewarded cue (CS-). Cue identity assignment was counterbalanced across animals such that half of the mice received the vertical cue as CS+ and the horizontal cue as CS-, and the other half received the opposite assignment.

On each trial, either the CS+ or CS- cue appeared at one of three spatial locations: the center monitor, TT/5.5 radians to the left of center, or TT/5.5 radians to the right of center. These spatial offsets were chosen to produce clearly distinguishable visual locations within the mouse visual field while remaining within the monitor array. On CS+ trials, mice received a 7 µL water reward 3 s after cue onset. The cue remained on the screen for the entire outcome period to avoid neural signal confounds caused by sudden cue disappearance. The outcome period lasted 4 s and was followed by an inter-trial interval of 8-15 s. On CS- trials, the cue remained on the screen for the same duration, but no reward was delivered.

Cue identity and spatial location were pseudo-randomized, with no more than three consecutive trials sharing the same cue identity and spatial location. During the entire session, mice were free to run on the Styrofoam ball. Ball movement was recorded and stored for later movement analysis. However, mouse movement was not coupled to the visual scene, and cue position remained stationary on the monitors regardless of locomotion. Task contingencies and trial structure were controlled using custom MATLAB scripts.

Approximately five days before three-location Pavlovian training, mice were acclimated to the head-fixed setup for 10-30 min per day. During acclimation, unexpected water rewards (7 µL) were delivered at random intervals drawn from a uniform distribution between 5 and 30 s. Imaging was also performed during this period to habituate mice to the imaging setup. After acclimation, mice were trained on the three-location Pavlovian task for approximately 30 min per session while neural activity was simultaneously recorded. Once task training and imaging began, mice were trained and imaged continuously, with no more than 1-3 day breaks between sessions, until behavioral performance indicated that the task had been learned. Data from the entire learning period and at least 5-7 days of stable learned performance were collected for analysis.

### Photometry recording

Fiber bundle imaging was performed using a custom microscope mounted on a vibration isolation table (Newport). Excitation light was provided by high-power 405 nm and 470 nm LEDs (Thorlabs SOLIS series). Excitation light was filtered (Chroma ET405/10 and ET473/24), combined using a dichroic mirror (Chroma 405rdc), and coupled into a liquid light guide (Newport 77632) through a series of lenses and a collimating beam probe. The light guide was connected to a filter cube on the microscope, and excitation light was reflected into the back aperture of a 10× objective (0.3 NA, Olympus UPLFLN10X2) by a dichroic beam splitter. Fluorescence emitted from the fiber bundle was collected through the same objective, passed through the dichroic beam splitter, and filtered with a green emission bandpass filter (Chroma 525/50) to remove residual excitation light and autofluorescence. Emission light was then focused onto a CMOS camera (Hamamatsu ORCA-Fusion BT Gen III) using a tube lens (Thorlabs TTL165-A), forming an image of the proximal fiber bundle face. Imaging data were acquired using HCImage Live (Hamamatsu). TTL pulses were returned from the camera to the acquisition card after each frame exposure to confirm proper triggering and align imaging data with behavioral recordings.

Calcium and glutamate signals were recorded using 470 nm excitation. In most recordings, 470 nm excitation was delivered at 30 Hz. In a subset of recordings, dual-wavelength excitation was used to control for calcium or glutamate independent fluctuations, including motion artifacts, photobleaching, and hemodynamic changes. For dual-wavelength recordings, the 470 nm wavelength excited sensor-dependent fluorescence, whereas 405 nm excitation served as an isosbestic control wavelength. The 405 nm signal was used to estimate non-neural fluctuations and to correct the 470 nm signal during analysis. The 405 nm and 470 nm LEDs were triggered by 5 V digital TTL pulses and alternated at 18 Hz, with a 20 ms exposure per frame. LED trigger pulses were sent in parallel to the camera to synchronize excitation with image acquisition. The timing and duration of digital pulses were controlled by custom MATLAB software through a programmable data acquisition card (National Instruments PCIe-6343).

### Post-mortem fiber localization

At the end of behavioral training and neural recordings, mice were injected intraperitoneally with Euthasol (400 mg/kg; Covertus Euthanasia III) and perfused intracardially with phosphate-buffered saline (PBS; Fisher Scientific), followed by 4% paraformaldehyde in PBS. Mice were decapitated, and the ventral side of the skull was removed to expose the ventral surface of the brain. Intact implanted brains were post-fixed for 24 h in 4% paraformaldehyde, rinsed with PBS, and transferred to Lugol iodine solution (Carolina Scientific) diluted 1:3 in distilled water for 4-6 days. Lugol iodine enhanced tissue contrast for computed tomography imaging and enabled fiber localization. CT scanning and fiber localization were performed as described previously ^51^. Briefly, 3D CT scans of intact implanted brains were registered to the Allen Mouse Brain Common Coordinate Framework atlas using a semi-automated landmark-based approach. Fibers appeared bright in CT scans and were automatically identified by intensity thresholding. Recording locations, defined as the ventral-most point of each fiber, were mapped to their corresponding positions on the implanted grid and subsequently to their positions on the proximal imaging surface. Registration to the atlas also brought all recording locations into a common coordinate space and enabled automatic assignment of anatomical labels, which were manually verified.

### Quantification and statistical analyses

#### Analysis cohorts and inclusion

Calcium recordings were obtained from Drd1-Cre and Adora2a-Cre mice expressing jGCaMP7f in dSPNs and iSPNs, respectively. The full calcium cohort included 8 Drd1-Cre mice and 6 Adora2a-Cre mice, corresponding to 430 dSPN-targeted and 329 iSPN-targeted fiber locations after post hoc localization. One Drd1-Cre mouse was recorded only after learning, whereas two Drd1-Cre mice with pre-learning recordings did not meet the behavioral learning criterion. As a result, pre-learning and learned-stage calcium analyses each included 7 Drd1-Cre mice, but these were not identical sets of animals.

Analyses of pre-learning visual responses used recordings obtained before conditioning from 7 Drd1-Cre mice and 6 Adora2a-Cre mice. Learned-stage analyses of cue-location encoding were restricted to mice trained in the three-location Pavlovian task with sufficient lateralized cue trials, yielding 6 Drd1-Cre mice and 4 Adora2a-Cre mice. Learned-stage cue-value and ramping analyses included these mice plus one additional Drd1-Cre mouse and two additional Adora2a-Cre mice trained in a one-location cue-association task, because these analyses did not require comparisons across cue locations. Thus, after session and fiber inclusion criteria were applied, learned-stage value and ramping analyses included 7 Drd1-Cre mice and 6 Adora2a-Cre mice.

Glutamate recordings were obtained from separate Drd1-Cre and Adora2a-Cre cohorts expressing membrane-targeted iGluSnFR in dSPNs or iSPNs. Value- and ramping-related glutamate analyses included 3 Drd1-Cre mice and 4 Adora2a-Cre mice, corresponding to 215 dSPN-targeted and 242 iSPN-targeted sites. Cue-location encoding analyses included the same 3 Drd1-Cre mice but only 3 Adora2a-Cre mice, because one Adora2a-Cre mouse was not trained in the three-location task, yielding 185 iSPN-targeted sites for the location analysis. These glutamate datasets were analyzed separately for cue-location, cue-value, ramping, and lick-related signals, with analysis-specific site counts reported in the Results and figure legends because some analyses required lateralized trials, significant cue responses, or restriction to anatomically defined regions.

## Behavior

### Learning phase definition

When mice learned the cue-reward association, they typically exhibited anticipatory licking before reward delivery. Anticipatory licking was quantified on each trial as the mean lick probability during the final second before reward onset. Lick probability was quantified for all six trial types, defined by cue value and cue location. To assess the development of Pavlovian learning, lick probability across trial types was examined over the course of training. To test whether mice distinguished CS+ from CS- trials across different cue locations, lick probability was analyzed using a linear mixed-effects model with cue value (CS+ versus CS-), cue location (contralateral, ipsilateral, or middle), and their interaction as fixed effects, with random intercepts for mouse and session nested within mouse. The model was specified as:

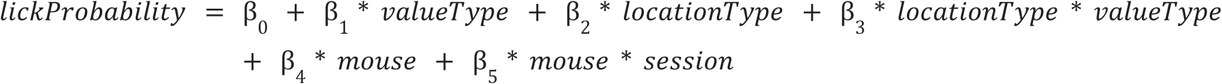

Successful discrimination between CS+ and CS- trials at each cue location was determined by post hoc comparisons of CS+ versus CS- within each location.

### Velocity

Velocity-related movement variables were extracted from spherical treadmill signals and expressed in a common implant-centered reference frame across animals. Angular velocity was computed from ball yaw velocity, converted to rad/s, and sign-flipped for left-implanted animals such that positive values consistently indicated contralateral movement and negative values indicated ipsilateral movement. Linear velocity was defined from pitch and roll ball velocities, converted to m/s, as:

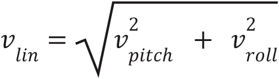

To capture both movement direction and magnitude, we defined additional movement variables, including directional velocity and change in directional velocity. Directional velocity was defined as the Euclidean magnitude of the three treadmill components, with its sign determined by yaw velocity:

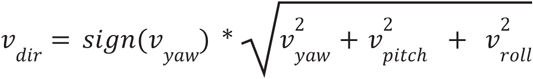

Under this definition, positive values indicate contralateral-directed movement, whereas negative values indicate ipsilateral-directed movement.

To reduce abrupt artifactual sign changes that can occur when linear velocity is high but yaw velocity fluctuates near zero, directional velocity was blended with the corresponding unsigned velocity magnitude using a yaw-dependent logistic weighting function:

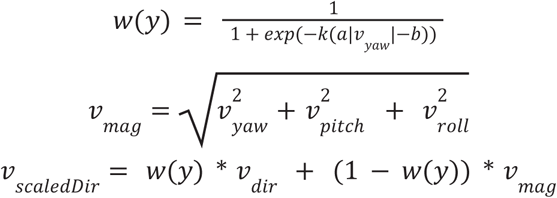

The parameters (k), (a), and (b) were chosen empirically based on model performance and were confirmed to yield stable results. In this formulation, the contribution of the signed directional component increases with the magnitude of yaw velocity, whereas the unsigned magnitude dominates when yaw velocity is small. The resulting scaled velocity was used as the directional velocity measure in subsequent analyses.

All velocity traces, including linear velocity, angular velocity, and directional velocity, were preprocessed identically. Raw signals were low-pass filtered at 1.5 Hz and then smoothed with a moving-average window of approximately 0.3 s. Change in directional velocity was computed from the smoothed directional velocity trace using a finite difference over an approximately 0.5 s interval, followed by a second low-pass filter at 1.5 Hz and an additional moving-average smoothing step of approximately 0.3 s. Under this convention, positive values of change in directional velocity indicate contralateral acceleration or ipsilateral deceleration, whereas negative values indicate ipsilateral acceleration or contralateral deceleration. Thus, these variables capture signed changes in movement along the contra-ipsi axis.

### Licking

Licking behavior was detected from the raw spout-contact signal. For analyses requiring a continuous licking regressor, the lick signal was converted to a time-resolved lick probability or lick-rate trace and processed using the same filtering and smoothing procedures described above. Lick initiation was defined on a trial-by-trial basis as the first time point at which the raw spout-contact signal exceeded a session-specific threshold and remained continuously above threshold for approximately 0.5 s.

### Movement-matched trial selection

For the analyses shown in Supplementary Fig. 3A-G and Supplementary Fig. 6A-D, we selected subsets of trials with matched movement profiles across conditions. For cue-location analyses, trials were matched between contralateral and ipsilateral cue conditions. For cue-value analyses, trials were matched between CS+ and CS- conditions.

To identify movement-matched trial pairs, cue-evoked changes in linear velocity, angular velocity, and directional velocity were quantified for each trial as the signed maximal deviation from baseline within the post-cue analysis window. Trials from the two conditions were then paired one-to-one by minimizing the absolute difference in directional velocity. Candidate matched subsets were ranked based on pairwise directional-velocity differences, and the largest subset was selected such that paired Wilcoxon signed-rank tests detected no significant difference between conditions in either linear velocity or angular velocity (p > 0.05; minimum 10 pairs).

### Neural activity (photometry)

#### Multifiber photometry preprocessing

Time-series movies were first motion-corrected using a whole-frame cross-correlation algorithm described previously^51^. Corrected movies were visually inspected to verify image stability, and recordings with excessive motion artifacts were excluded from further analysis. Mean fluorescence signals were extracted from circular regions of interest (ROIs) corresponding to individual fibers. ROI radii were set to approximately half the fiber diameter, 25 µm for 50 µm fibers and 19 µm for 37 µm fibers. The same ROI map was applied across recordings within each experiment, with minor adjustments to accommodate shifts in the imaging field and ensure consistent fiber identification across sessions. Relative fluorescence changes (.LF/F) were calculated by normalizing extracted fluorescence traces to a baseline defined as the 8th percentile of fluorescence values within a 30 s sliding window. This procedure reduced slow baseline fluctuations arising from photobleaching or other low-frequency signal drift.

#### Identification and temporal characterization of cue-evoked responses

Cue responses were quantified from z-scored fluorescence signals for each fiber ROI. Fluorescence traces were z-scored within each fiber before trial averaging. Trials of the relevant condition were selected based on trial labels. For each fiber, activity was averaged across trials and analyzed in a 0-3 s window after cue onset, spanning the cue period. The signal was baseline-corrected by subtracting the mean activity during the 1 s pre-cue baseline. Cue responsiveness was determined using a bootstrap-based threshold. A null distribution was generated by sampling pseudo-events from non-event periods of the recording while excluding real events and other excluded intervals. For each bootstrap iteration (n = 5000), pseudo-events equal in number to the real events were sampled with replacement. Event-aligned activity was extracted and baseline-corrected using the same procedure as for the real events, and activity was averaged across sampled pseudo-events. This procedure generated a time-resolved null distribution of mean activity for each fiber.

Significance thresholds were defined from the bootstrap distribution. A fiber was classified as cue responsive if its baseline-corrected mean signal exceeded the 99.5th percentile of the bootstrap distribution for at least five consecutive samples within the 0-3 s post-cue window. For responsive fibers, peak latency was defined as the time of the first local maximum within the significant response segment. Local maxima were detected using a minimum prominence threshold equal to 30% of the segment’s maximum amplitude. If no peak satisfied this criterion, latency was assigned as the time of the maximum signal within the significant segment.

#### FIR-based elastic net regression for quantifying cue-location and cue-value encoding

To quantify cue-location and cue-value encoding at each recording site, z-scored neural activity traces were concatenated across included trials and modeled as a linear combination of finite impulse response (FIR) kernels aligned to cue onset. The model included a cue-onset kernel, a task-variable difference kernel, and movement-related covariates. For cue-location analyses, only lateralized trials were included, and the task variable encoded cue location relative to the implanted hemisphere, with ipsilateral trials assigned *x^task^* = 0 and contralateral trials assigned *x^task^* = 1. For cue-value analyses, the task variable encoded cue value, with CS- trials assigned *x^task^* = 0 and CS+ trials assigned *x^task^* = 1. Thus, the task-variable kernel captured either the contralateral-minus-ipsilateral difference in cue-evoked activity or the CS+-minus-CS- difference in cue-evoked activity. Positive location-kernel coefficients indicate stronger responses on contralateral trials, and negative coefficients indicate stronger responses on ipsilateral trials. Positive value-kernel coefficients indicate stronger responses on CS+ trials, whereas negative coefficients indicate stronger responses on CS- trials.

In analyses testing whether cue-location or cue-value encoding was explained by licking, trials with early licking were excluded by removing trials in which lick initiation occurred within approximately 1.5 s of cue onset. Lick initiation criteria are described above. In analyses testing whether cue-location or cue-value encoding was explained by movement, movement-matched trial subsets were used as described above.

The model was fit over a time window spanning from 1 s before to 1.5 s after cue onset. Time-resolved coefficients were estimated using elastic net regression in MATLAB:

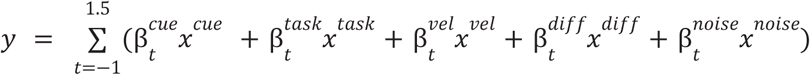

where *t* spans from -1 s to 1.5 s relative to cue onset. The cue regressor, *x^cue^*, was set to 1 for all included cue trials. Under this parameterization, 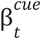 estimates the predicted response to the reference condition, corresponding to ipsilateral trials for cue-location analyses and CS- trials for cue-value analyses. The sum of 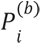 and 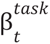 estimates the predicted response to the comparison condition, corresponding to contralateral trials for cue-location analyses and CS+ trials for cue-value analyses.

Movement-related covariates were included to account for variance over the same peri-cue interval. Specifically, *x^vel^* represents directional velocity, and *x^diff^* represents change in directional velocity as described above. A Gaussian noise term, *x^noise^*, was included as a control regressor. Models were fit separately for each ROI. Hyperparameters for elastic net regression were selected by grid search over a and ’J, and model performance was evaluated using 5-fold cross-validation. Coefficients were baseline-corrected by subtracting the mean value during the 1 s pre-cue period. This approach quantified cue-location or cue-value encoding as a difference in cue-evoked activity between task conditions while controlling for concurrent movement-related signals.

#### Location-Value interaction analysis

To test whether cue-value encoding depended on cue location at individual recording sites, we restricted the analysis to sites with significant encoding in both the cue-location and cue-value analyses described above. Using only lateralized trials, baseline-corrected z-scored activity was measured at the peak latency identified in the cue-value analysis. A linear model was then fit separately for each recording site:

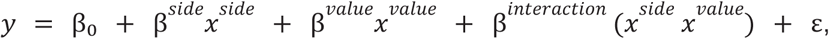

where *x^side^* encoded cue location with ipsilateral trials relative to the implanted hemisphere assigned 0 and contralateral trials assigned 1. Cue value was encoded as *x^value^* = 1 for CS+ trials and *x^value^* = 0 for CS- trials. Under this parameterization, β^*interaction*^ captures the extent to which cue-value encoding differed between contralateral and ipsilateral cue locations.

#### Orthogonal Time-Varying Basis Models for Identifying Ramping Activity

To characterize ramp-like temporal structure in trial-resolved fiber photometry signals, activity within the analysis window was modeled using a small set of orthogonal time-varying basis functions. For each trial, activity from each fiber was extracted from 0 to 3 s relative to cue onset, baseline-corrected using the 1 s pre-cue period, and standardized within the modeled window. This preprocessing reduced the influence of between-trial differences in absolute amplitude and emphasized the temporal profile of the signal. This yielded a discrete orthonormal set of time bases, denoted P_0_, P_1_, …, P_k_, where P_0_ corresponds to a constant component, P_1_ represents a linear temporal trend, and higher-order terms capture progressively more complex nonlinear temporal structure. Because the basis functions were mutually orthogonal, the coefficient associated with each term could be interpreted as the strength of a distinct temporal component with reduced collinearity among regressors.

##### Basic model

In the basic model, the standardized activity trace from each trial was expressed as a linear combination of these orthogonal time bases:

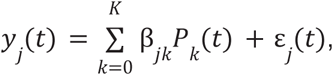

where *y*_*j*_(*t*) is the signal from trial *j*, β_*jk*_ is the coefficient for basis *P_k_*, and £_*j*_(*t*) is the residual error. The coefficient of *P*_1_ was treated as the primary index of linear ramping. For each fiber, coefficients were estimated separately for each trial, and a right-tailed t-test was used to assess whether the across-trial distribution of *P*_1_ coefficients was greater than zero. Fibers with significant positive *P*_1_ coefficients were classified as ramping sites. This formulation decomposed temporal dynamics into constant, linear, and higher-order nonlinear components, allowing the linear term to serve as an interpretable index of ramping.

##### Basic model restricted to pre-lick data

To determine whether ramping activity was present before lick initiation, we implemented a pre-lick variant of the same model. This analysis was restricted to trials in which lick onset occurred more than 1 s after cue onset. For each retained trial, only fluorescence signals preceding the trial-specific lick onset were included. Because the duration of the pre-lick interval varied across trials, the orthogonal time basis was reconstructed separately for each trial segment and used to fit the pre-lick fluorescence signal.

##### Basic model with lick-rate covariate

To further dissociate ramp-like temporal structure from movement-related activity, lick rate was added as an additional trial-wise regressor to the model. Lick rate was baseline-corrected and standardized in the same window as the neural signal. The resulting model was

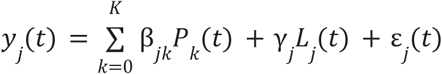

where *L_j_*(*t*) is the lick-rate trace for trial *j*, and *y_j_* is its regression coefficient. In this formulation, *P*_0_ served as the constant term, so no additional intercept was included. The coefficient of *P*_1_ in this model quantified the linear ramp component after accounting for trial-by-trial variation in licking.

#### Prediction of lick initiation latency by ramp slope

For each animal, we asked whether trial-by-trial ramping activity predicted lick initiation latency using repeated 5-fold cross-validated linear regression. For each selected fiber, trial-by-trial ramp slope was estimated from the fluorescence signal using the orthogonal time-varying basis model described above. Lick initiation latency was defined as the first time point at which licking began and persisted for at least 0.5 s. Only fibers with significant ramping activity were included in this analysis.

For each animal, ramp slopes from all selected fibers were combined into a trial-by-fiber predictor matrix and used to predict lick initiation latency:

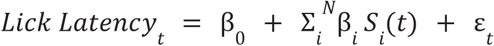

where *S_i_* (*t*) is the ramping slope of fiber i on trial t, and *N* is the number of ramping fibers included for that animal.

To evaluate predictive performance, trials were randomly divided into five folds. For each fold, the model was fit using 80% of trials and tested on the held-out 20%, yielding a cross-validated R^2^. This 5-fold cross-validation procedure was repeated 50 times with independent fold assignments for each animal, and the mean cross-validated R ^2^ across repeats was used as the observed prediction performance.

To test whether prediction exceeded chance, lick initiation latencies were randomly shuffled across trials while the ramping slope matrix was kept unchanged. The same repeated 5-fold cross-validation procedure was applied to each shuffled dataset, and the mean shuffled R^2^ across 50 repeats was used as one null sample. This label-shuffle procedure was repeated 500 times to generate a null distribution of shuffled mean R^2^ values. Statistical significance was assessed using a one-sided empirical p value, computed as the fraction of shuffled mean R^2^ values greater than or equal to the observed mean R^2^, with a +1 correction:

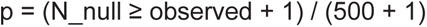

### Volumetric Analyses

#### Rasterization and smoothing

For volumetric analyses, site-specific data were rasterized onto a three-dimensional voxel grid spanning -2.6 to 1.9 mm along the anterior-posterior axis, 0.3 to 4.1 mm along the medial-lateral axis, and -6.0 to -1.5 mm along the dorsal-ventral axis. The voxel size was 0.05 mm. A striatal mask was applied to include only voxels located within the striatum.

Smoothed fiber measurements, including coefficient amplitudes and raw ΔF/F values, were estimated on this three-dimensional striatal voxel grid using a spherical neighborhood and a Laplace distance kernel. Specifically, the value at each voxel was computed as a normalized weighted sum of amplitudes from fibers located within a radius *r*, with weights defined as:

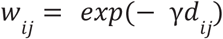

for fibers within the neighborhood and 0 otherwise. Here, *i* denotes the target voxel in the striatal volume, *j* denotes an individual fiber, A_j_ is the measurement amplitude of fiber *j*, and *d_ij_* is the Euclidean distance between voxel *i* and the location assigned to fiber *j*.

The parameters y and *r* were selected empirically based on model performance and were confirmed to yield stable results.

The final estimate at each voxel was computed as:

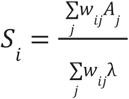

where ’J is an adaptive regularization term defined as the 0.2 quantile of the voxel-wise total kernel weight distribution. This term was introduced to mitigate inflated or unstable estimates in voxels with sparse fiber coverage.

Horizontal and sagittal views were generated by mean-projecting the smoothed 3D volume into two dimensions. Horizontal maps were obtained by averaging nonzero voxel values across the dorsal-ventral axis at each anterior-posterior and medial-lateral coordinate. Sagittal maps were obtained by averaging nonzero voxel values across the medial-lateral axis at each anterior-posterior and dorsal-ventral coordinate.

#### Permutation-based estimation of significant threshold and contour identification

To reduce the influence of amplitude variability across fibers, significant contour were generated by smoothing the binary significant-fiber mask rather than the continuous fiber amplitude. Let *M_i_ E* {0. 1} denote the significance label of fiber *j*, where *M_i_* = 1 indicates a significant fiber and *M_i_* = 0 otherwise. For each voxel *i*, a smoothed significance value was computed using the same spherical neighborhood and Laplace distance kernel used for volumetric smoothing:

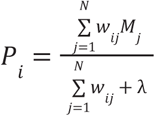

where *i* indexes voxels, *j* indexes fibers, *N* is the total number of fibers, w_ij_ = exp(- yd_ij_) for fibers located within radius *r* of voxel *i* and w_ij_ = 0 otherwise, d_ij_ is the Euclidean distance between voxel *i* and the voxel assigned to fiber *j*, and ’J is an adaptive regularization term.

Under this formulation, P_i_ can be interpreted as a local probability-like estimate that voxel *i* is associated with significant fibers.

To estimate the null distribution, the significant-fiber labels were permuted across fiber identities while preserving the total number of significant fibers. For each permutation *b*, the shuffled significant mask was spatially smoothed using the same kernel, radius, and normalization parameters, yielding a permuted voxelwise map 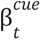. The 95th percentile of the voxelwise values from each permuted map was then recorded:

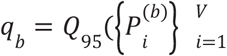

where *V* is the total number of voxels in the striatal grid. After 1000 permutations, the significance threshold was defined as the mean of these permutation-specific 95th percentile values.

The observed smoothed significance map was then thresholded against this permutation-derived probability threshold, indicating that voxel *i* exceeded the null expectation for the local concentration of significant fibers.

For contour visualization, the resulting binary significance volume was further regularized by 3-dimensional box smoothing with a uniform kernel, thresholded at 0.2, and partitioned into connected components using 26-neighbor connectivity. The boundaries of the retained components were then extracted and plotted as the significant contour in the horizontal and sagittal views.

#### Bootstrap-based assessment of spatial consistency across animals

To assess the spatial consistency of encoding across animals, we asked whether signals encoding value or location were preferentially concentrated within the anatomically defined value-encoding region in each individual mouse. For each mouse, fibers located inside this region were identified and their encoding strength was compared with a within-mouse null distribution generated by repeatedly sampling the same number of fibers from the full recorded population. This analysis allowed us to determine whether the observed value or location-related activity within the region was greater than expected by chance given that animal’s overall sampling distribution.

#### Spatial overlap analysis

To assess whether two or more contours overlapped beyond chance levels, we quantified the observed overlap as the number of striatal voxels jointly contained within all contours under comparison. For analyses involving more than two contours, overlap was defined as the common intersection across all contours, rather than as the sum of pairwise overlaps.

Statistical significance was evaluated using an empirical null distribution. For each contour, surrogate volumes were generated by randomly placing spatially contiguous regions within the striatum while preserving the original contour size. For each iteration, the shared volume among the surrogate contours was computed, and this procedure was repeated to generate a distribution of overlap values expected by chance.

The observed overlap was then compared to this null distribution. An empirical one-sided P value was calculated as the fraction of surrogate overlaps that were equal to or greater than the observed overlap. Overlap was considered significant when it exceeded chance expectation at a threshold of p < 0.05.

#### Identification of convergent and divergent value-encoding contour

To identify striatal regions in which dSPNs and iSPNs exhibited either shared or opponent value encoding, we computed a voxelwise multiplication map from the volumetric value-encoding maps of the two cell types. For each voxel, the value-encoding metric from the dSPN map was multiplied by the corresponding metric from the iSPN map. Positive multiplication values indicated that dSPNs and iSPNs encoded value with the same sign, whereas negative multiplication values indicated opponent-signed value encoding between the two cell types.

Statistical significance was evaluated on a voxelwise basis using a permutation-based null distribution. Specifically, 10,000 null multiplication maps were generated by randomly pairing dSPN and iSPN permutation maps and recomputing the voxelwise product. For each voxel, a one-tailed p-value was calculated by comparing the observed multiplication value with the corresponding null distribution: for positive observed values, p was defined as the fraction of null values greater than or equal to the observed value; for negative observed values, p was defined as the fraction of null values less than or equal to the observed value.

Voxels were thresholded at p < 0.05, and spatially contiguous significant voxels were grouped into connected three-dimensional clusters. Significant contours were defined as connected volumes larger than 1077 voxels. This minimum cluster-size threshold was chosen to exclude small isolated clusters and was approximately matched to the voxel volume expected from a single outlier recording site after spatial smoothing, based on the estimated collection volume of an individual fiber^51^ and the Laplace smoothing procedure described above. Using this approach, positive contours identified regions with shared positive value encoding in dSPNs and iSPNs, whereas negative contours identified regions with opponent value encoding across the two cell types.

## Acknowledgements

This work was supported by a Klingenstein-Simons’s Foundation fellowship, Whitehall Foundation Fellowship, and National Institute of Mental Health (R01 MH125835) to M.W.H. We thank the Boston University Centers for Neurophotonics and Systems Neuroscience for financial and technical support and Boston University Animal Science Center for providing central laboratory and animal care and support resources. We thank Aaquib Attarwala for important pilot experiments that helped clarify the present work, Sydney Holder for assistance with CT scanning, and Chenxi Shi for optimizing code used in the volumetric analyses.

---

**Supplementary Figure 1.**
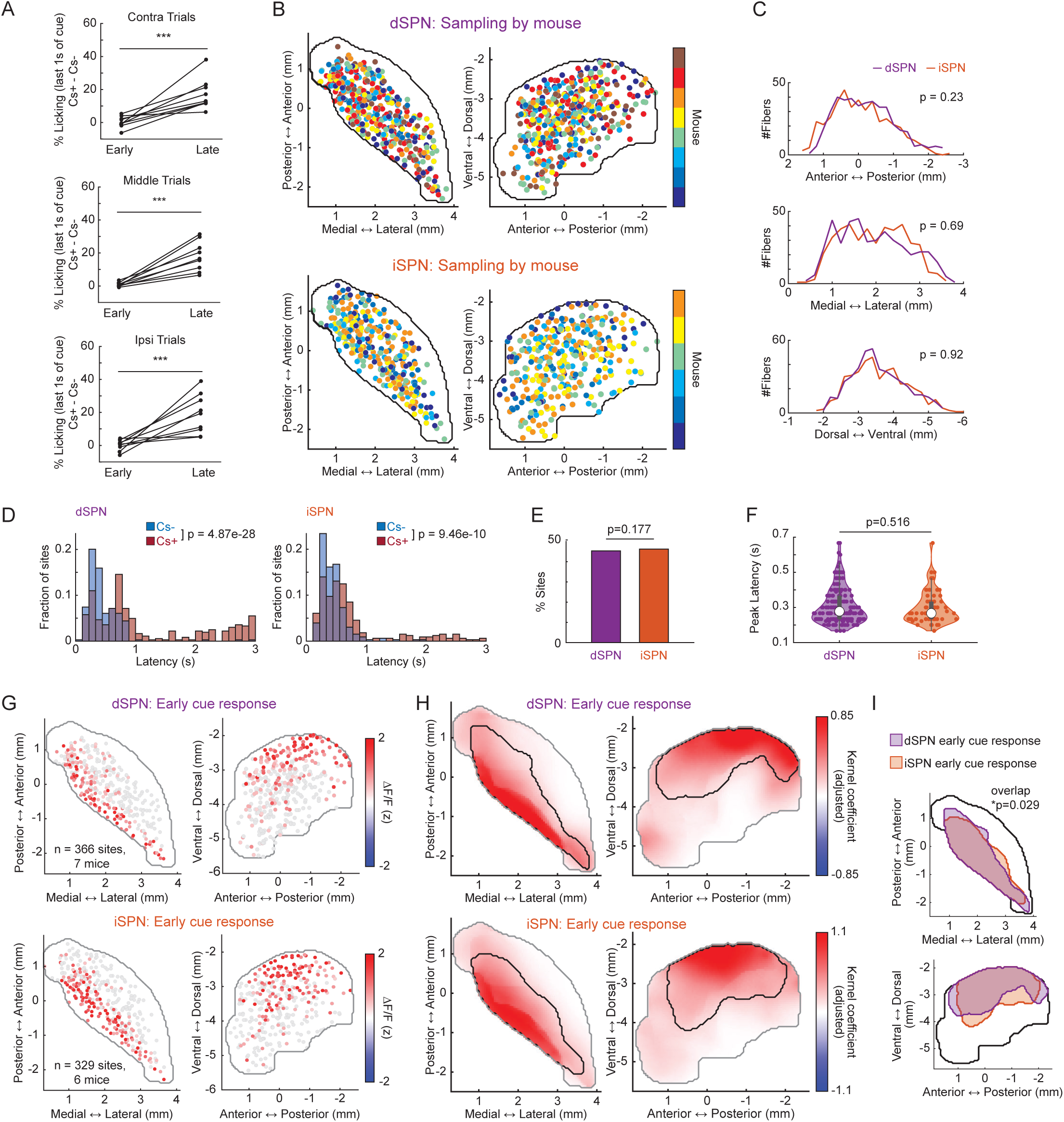
Calcium recording coverage and pre-learning cue responses (A) Cue discrimination increased across learning for contralateral, middle, and ipsilateral cue locations, defined relative to the implanted hemisphere. Each line represents one mouse and connects early and late learning values. Cue discrimination was quantified as CS+ minus CS-lick probability during the final second before reward delivery. Learning-related cue discrimination was assessed using a linear mixed-effects model of lick probability as a function of task stage, cue type, and their interaction, with random intercepts for mouse and session. Statistical significance was determined from the p value of the task stage × cue type interaction term. (B) Horizontal and sagittal distributions of striatal recording sites from dSPN and iSPN calcium-imaging cohorts. Each circle indicates one fiber location, and circle color denotes the individual mouse from which that recording site was obtained. (C) Sampling coverage along the anterior-posterior, medial-lateral, and dorsal-ventral axes. Purple and orange lines indicate the number of dSPN and iSPN calcium recording sites, respectively, at each coordinate value. Coverage along each axis was comparable between dSPN and iSPN cohorts. Two-sample Kolmogorov-Smirnov tests, anterior-posterior: p = 0.23; medial-lateral: p = 0.69; dorsal-ventral: p = 0.92. (D) Distributions of CS+ and CS- peak response latencies for dSPN and iSPN sites in the learned phase. CS- responses peaked at shorter latencies than CS+ responses in both cell types. Wilcoxon rank-sum tests, dSPN: p = 4.87 × 10 ^-28^; iSPN: p = 9.46 × 10^-10^. (E) Proportion of sites with significant early cue responses before learning. The fraction of early cue-responsive sites did not differ between dSPN and iSPN datasets. Chi-square test of independence, p = 0.177. (F) Distribution of pre-learning early cue-response peak latencies for dSPN and iSPN sites. Dots indicate individual recording sites. Latencies did not differ between cell types. Linear mixed-effects model with latency as a function of cell type and mouse as a random effect; p = 0.516. (G) Horizontal and sagittal maps of mean pre-learning early cue-evoked responses in dSPN (top) and iSPN (bottom) sites. Each circle indicates one recording site. Red intensity indicates cue-evoked activity strength, and gray circles indicate sites without significant cue-evoked activity. (H) Smoothed horizontal and sagittal maps of pre-learning early cue responses in dSPN and iSPN sites. Voxel color indicates cue-response strength after spatial smoothing. Black contours mark significant group-level early cue-response regions. (I) Overlay of significant pre-learning early cue-response regions for dSPN and iSPN datasets. The two regions significantly overlapped (p = 0.029). Spatial overlap was assessed using the spatial overlap test described in Methods.

**Supplementary Figure 2.**
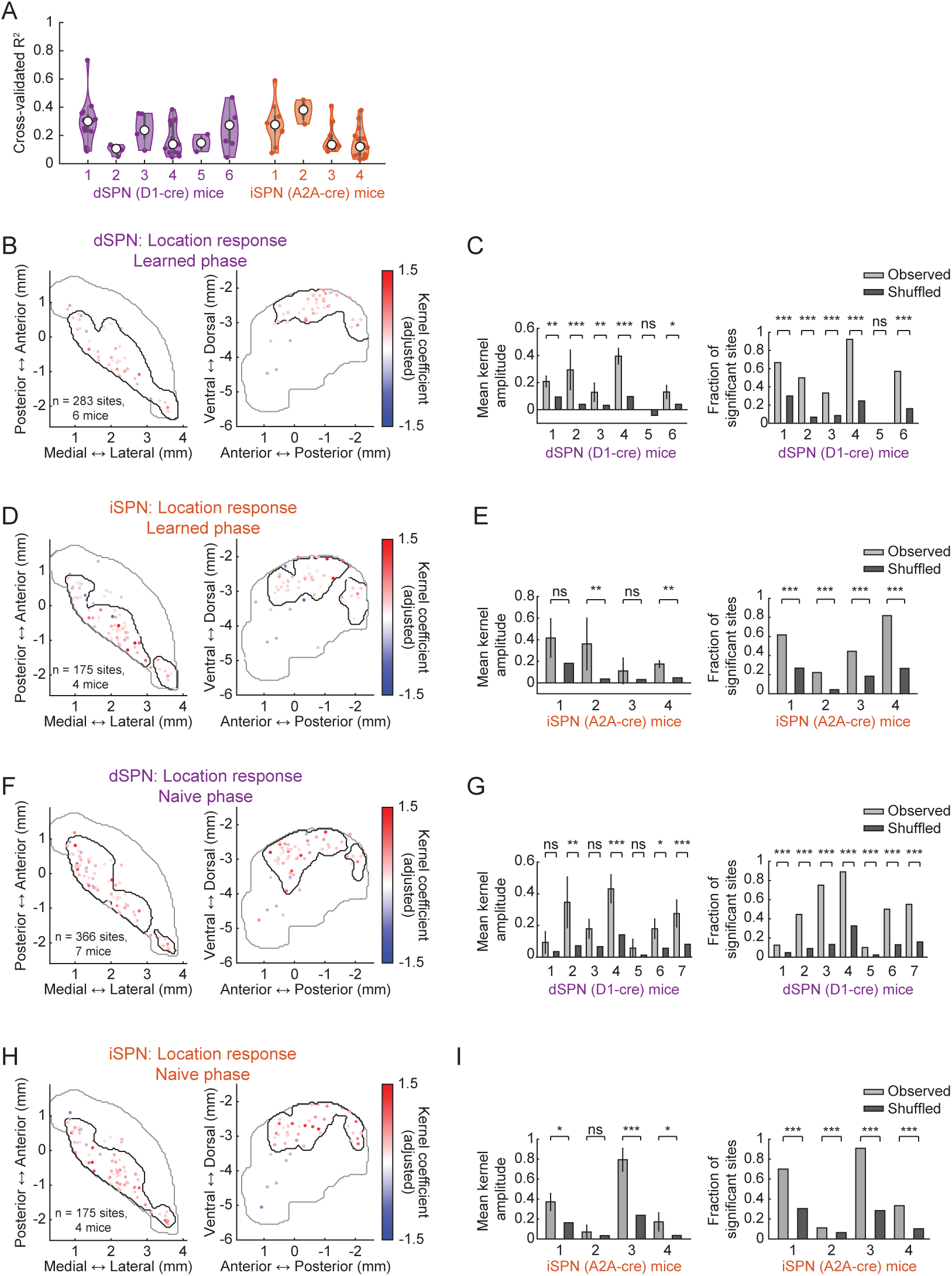
Model performance and spatial consistency of cue-location encoding (A) Cross-validated model performance for dSPN and iSPN datasets across mice. Violin plots summarize model fit, and dots indicate individual recording sites. Only sites with significant location encoding were presented. (B) Horizontal and sagittal maps of significant location-selective dSPN recording sites during the learned phase. Each dot indicates one significant site and is colored by location-kernel amplitude. Black contours mark the group-level significant location-encoding region. (C) Mouse-level quantification of learned-stage dSPN location encoding. Left, mean location-kernel amplitude for significant sites within the defined encoding contour in each mouse. Right, fraction of significant sites within the defined encoding contour in each mouse. Light gray bars show observed values; dark gray bars show shuffled controls generated by randomly selecting the same number of significant sites from across the striatum, independent of anatomical location. Statistical significance was assessed for each mouse by comparing observed contour values with the shuffled distribution using a linear mixed-effects model with shuffle groupID as a random effect. (D) Same as (B), for learned-stage iSPN sites. (E) Same as (C), for learned-stage iSPN sites. (F) Same as (B), for naive-stage dSPN sites. (G) Same as (C), for naive-stage dSPN sites. (H) Same as (B), for naive-stage iSPN sites. (I) Same as (C), for naive-stage iSPN sites. For learned-stage datasets, dSPNs include n = 283 sites from 6 mice and iSPNs include n = 175 sites from 4 mice. For naive-stage datasets, dSPNs include n = 366 sites from 7 mice and iSPNs include n = 175 sites from 4 mice. Asterisks indicate observed-versus-shuffled comparisons: *p < 0.05, **p < 0.01, ***p < 0.001; ns, not significant.

**Supplementary Figure 3.**
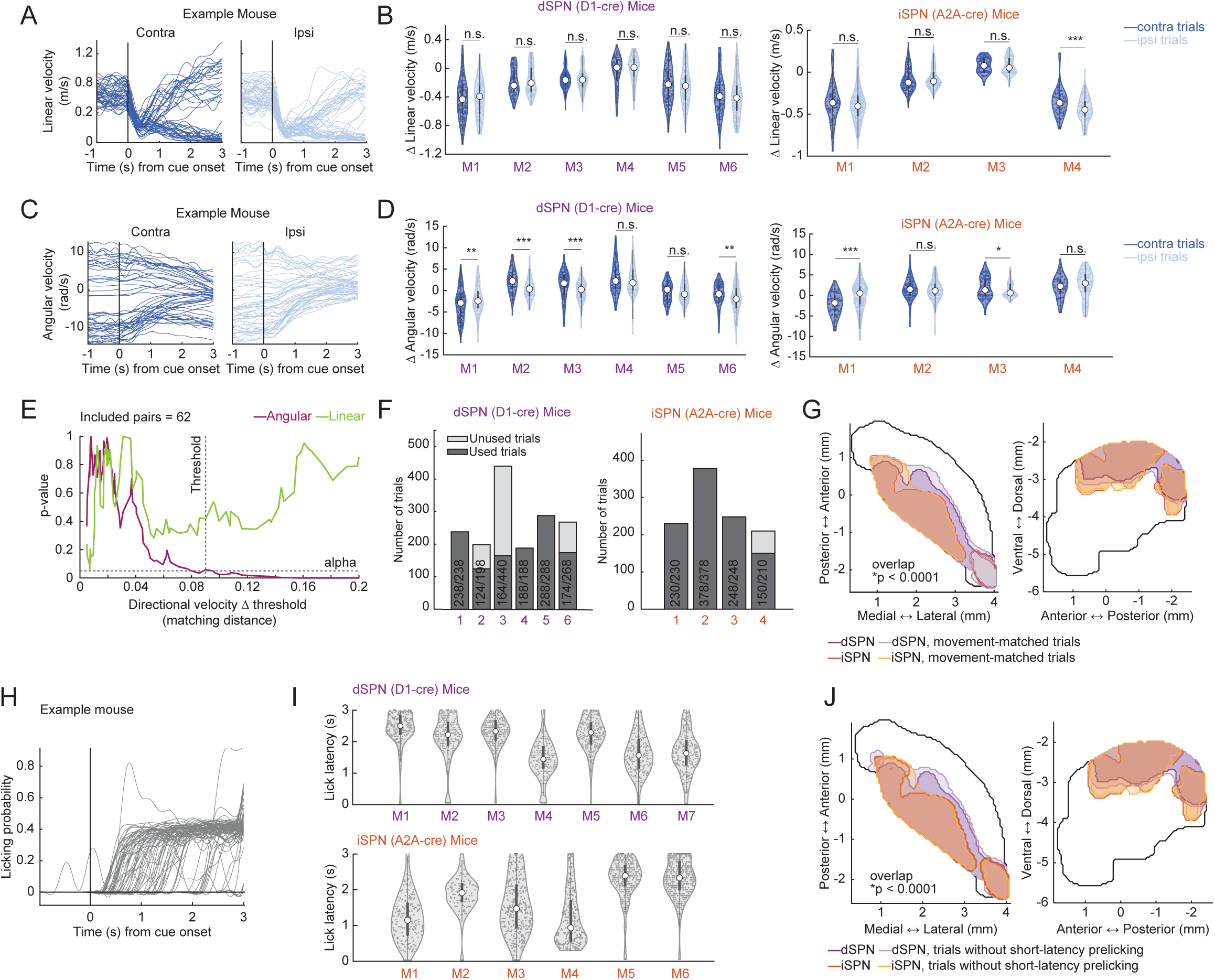
Locomotion and licking do not explain cue-location encoding (A) Example linear velocity responses for contralateral and ipsilateral cue trials in one mouse. Thin traces indicate individual trials. (B) Cue-evoked changes in linear velocity for contralateral and ipsilateral trials across dSPN and iSPN mice, quantified as the maximum change in linear velocity within 1 s after cue onset for each trial. Each dot represents one trial. Asterisks indicate significant contralateral-versus-ipsilateral comparisons, assessed by two-sample t-tests. (C) Same as (A), for angular velocity. (D) Same as (B), for angular velocity. (E) Example movement-matching procedure for selecting contralateral and ipsilateral trial pairs with matched velocity profiles. For each trial, cue-evoked changes in linear velocity, angular velocity, and directional velocity were quantified as signed maximal deviations from baseline within the post-cue analysis window. Trials were paired across contralateral and ipsilateral conditions by minimizing the absolute difference in directional velocity. Candidate subsets were then evaluated by paired Wilcoxon signed-rank tests comparing linear and angular velocity between conditions. Green and magenta traces show the p values for linear and angular velocity comparisons as a function of the directional velocity-difference threshold. The dashed horizontal line indicates the criterion for matched behavior, p > 0.05, and the dashed vertical line indicates the selected threshold at which neither linear nor angular velocity differed significantly between conditions. (F) Number of trials retained or excluded for movement-matched analyses in each dSPN and iSPN mouse. Dark bars indicate trials included in the movement-matched dataset, and light bars indicate unused trials. (G) Overlay of significant location-encoding contours from the full dataset and movement-matched trial subset for dSPN and iSPN recordings. Location-encoding regions derived from movement-matched trials significantly overlapped with those derived from all trials, indicating that pDMS cue-location encoding was preserved after controlling for contra/ipsi movement differences (p < 0.0001). (H) Example prelicking behavior during the cue period in one mouse. Each line indicates one trial. (I) Lick latency distributions for dSPN and iSPN mice. Each dot represents one trial. (J) Overlay of significant location-encoding contours from the full dataset and from trials without short-latency prelicking during the cue period. Location-encoding regions significantly overlapped across analyses (p < 0.0001). Spatial overlap was assessed using the spatial overlap test described in Methods. Asterisks indicate contralateral versus ipsilateral comparisons: *p < 0.05, **p < 0.01, ***p < 0.001; ns, not significant.

**Supplementary Figure 4.**
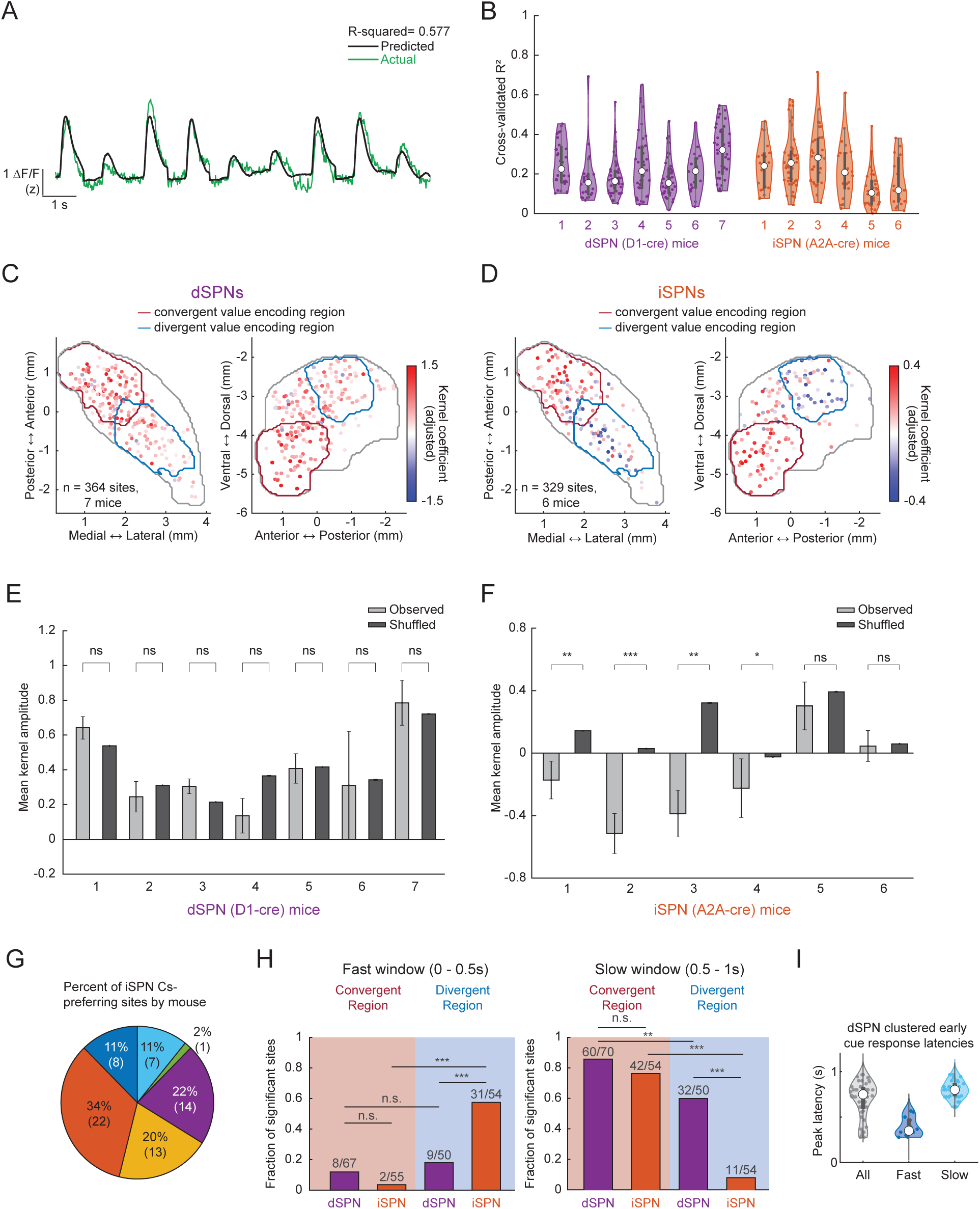
Model performance and spatial organization of cue-value encoding (A) Example model fit for the value-encoding model. Green trace shows the measured signal and black trace shows the model prediction. (B) Cross-validated model performance for dSPN and iSPN datasets across mice. Violin plots summarize model fit, and dots indicate individual recording sites. (C) Horizontal and sagittal maps of significant value-encoding dSPN sites. Each dot indicates one significant site and is colored by value-kernel amplitude. Red and blue contours denote the convergent and divergent value-encoding regions, respectively. (D) Same as (C), for iSPN sites. (E) Mouse-level quantification of dSPN value encoding. Bars show mean value-kernel amplitude for significant sites in each mouse. Light gray bars show observed values and dark gray bars show shuffled controls. (F) Same as (E), for iSPN sites. (G) Mouse composition of iSPN CS- preferring sites. Pie slices indicate the percentage and count of CS- preferring iSPN sites contributed by each mouse. All iSPN mice had at least one CS- preferring site. (H) Fraction of significant value-encoding sites in the fast window, 0-0.5 s after cue onset, and slow window, 0.5-1.0 s after cue onset, for dSPN and iSPN sites in the convergent and divergent value-encoding regions. Numbers above bars indicate significant value-encoding sites over total cue-responsive sites during the analyzed time window. Linear mixed-effects model with significance status predicted by encoding region and cell type and mouse as a random effect. (I) dSPN value-encoding latencies in the divergent region. Gray shows all sites, dark blue shows fast sites, and light blue shows slow sites after k-means clustering. Dots indicate individual recording sites. Asterisks indicate observed versus shuffled comparisons or model-based pairwise comparisons, as appropriate: *p < 0.05, **p < 0.01, ***p < 0.001; ns, not significant.

**Supplementary Figure 5.**
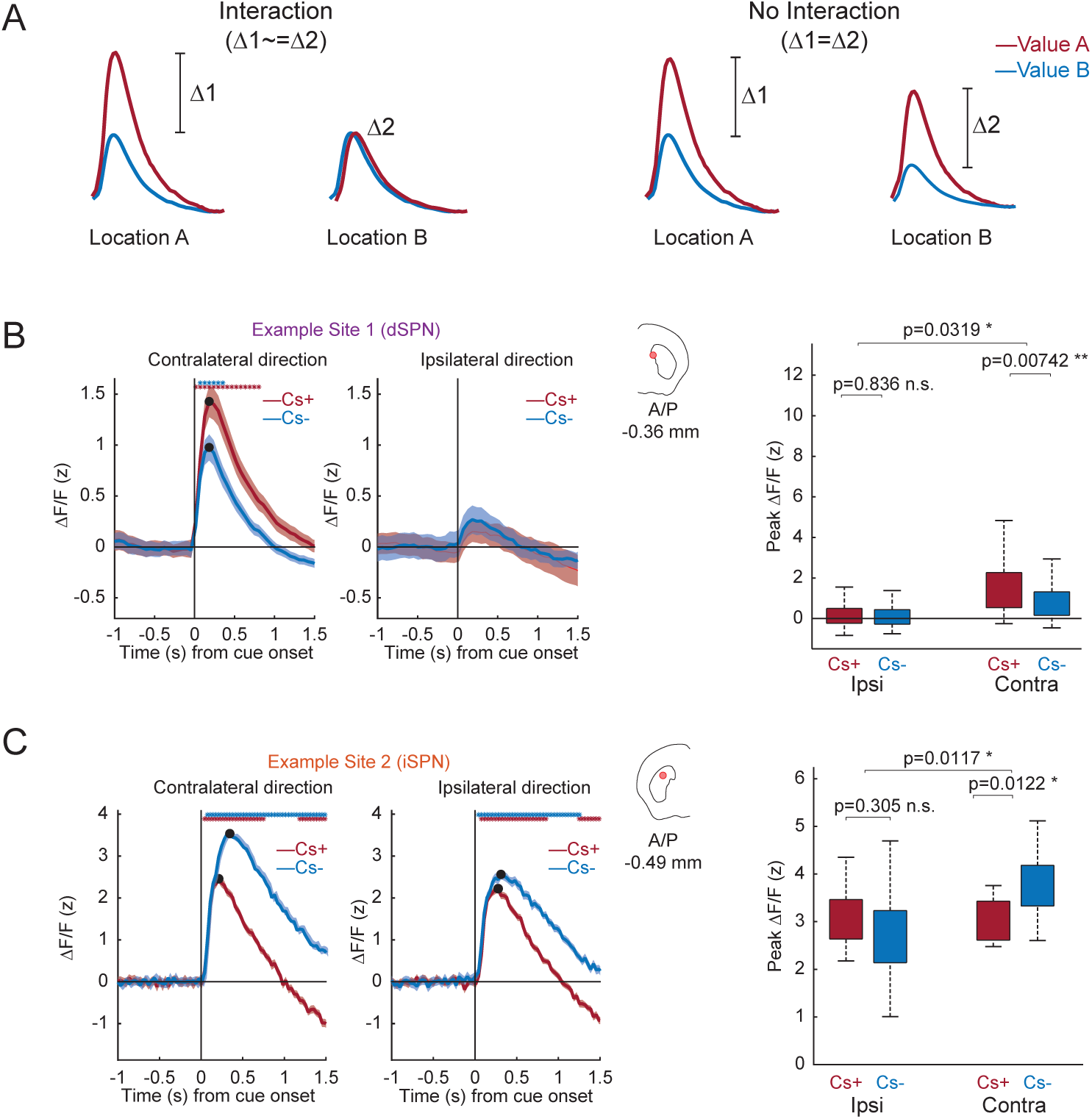
Value-location interactions in pDMS (A) Schematic illustrating independent versus interactive encoding of cue value and cue location. In the no-interaction case, value-related differences are similar across cue locations. In the interaction case, value-related differences depend on cue location. (B) Example dSPN site with a contralateral CS+ interaction. Left, mean responses to CS+ and CS- cues presented in contralateral and ipsilateral locations. Shaded regions indicate SEM. Coronal schematic indicates site location. Right, peak response amplitudes across cue-value and cue-location conditions. Box plots show trial-level distributions. Significance was assessed using a linear mixed-effects model with peak response as a function of location type, value type, and their interaction, with mouse included as a random intercept. Pairwise comparisons showed no significant difference between ipsilateral CS+ and CS- responses (p = 0.836), but a significant difference between contralateral CS+ and CS- responses (p = 0.00742). The location × value interaction was also significant (p = 0.0319). (C) Same as (B), an example iSPN site with a contralateral CS- interaction. Asterisks indicate post hoc comparisons: *p < 0.05, **p < 0.01; ns, not significant.

**Supplementary Figure 6.**
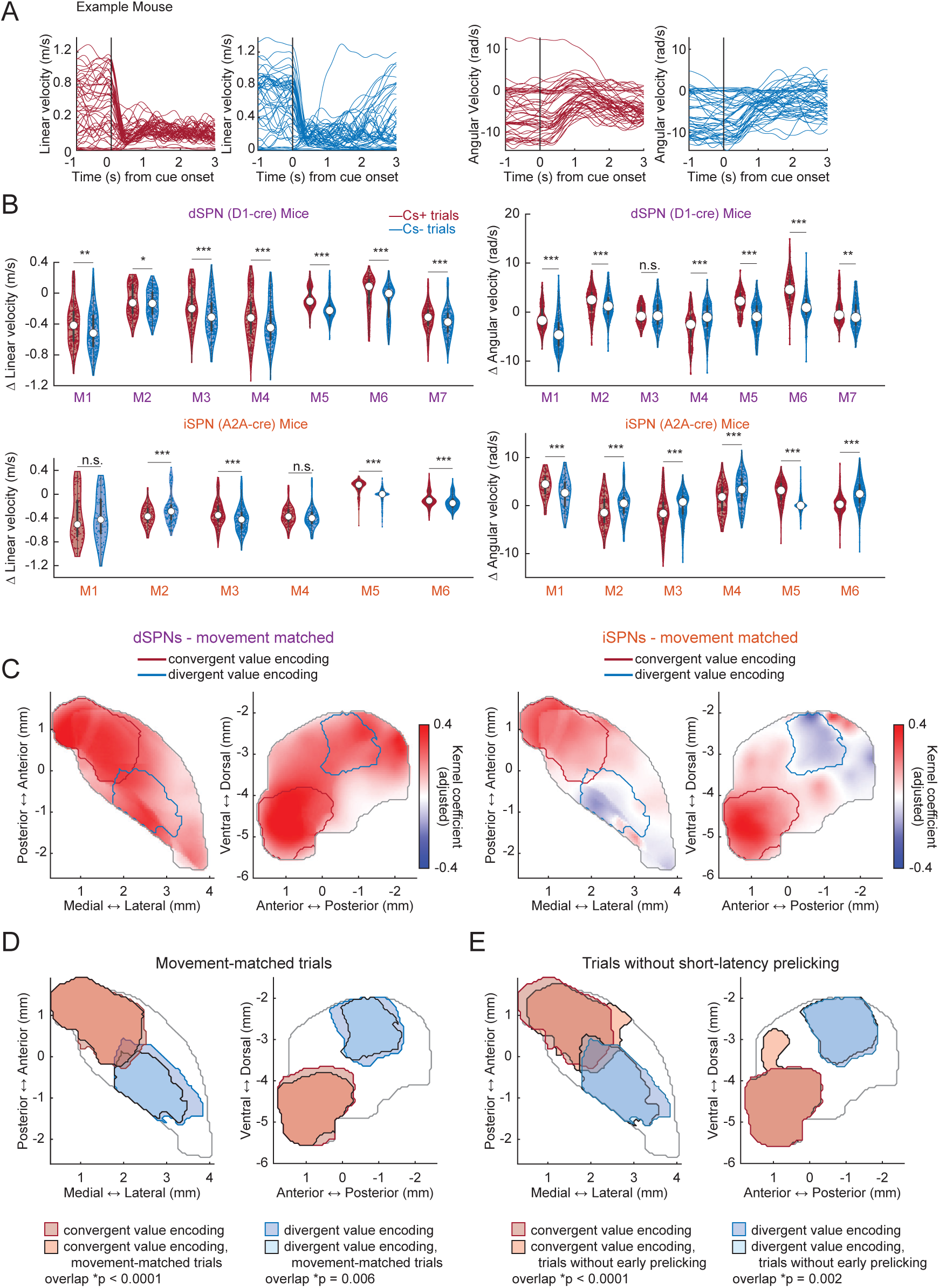
Locomotion and licking do not explain cue-value encoding (A) Example linear velocity and angular velocity responses for CS+ and CS- trials in one mouse. Thin traces indicate individual trials. Red traces show CS+ trials and blue traces show CS- trials. (B) Cue-evoked changes in linear and angular velocity for CS+ and CS- trials across dSPN and iSPN mice, measured as the maximum velocity change within 1 s after cue onset. Each dot represents one trial. Asterisks indicate significant CS+ versus CS- comparisons, assessed by two-sample t-tests. (C) Smoothed horizontal and sagittal maps of value encoding after movement matching for dSPN and iSPN datasets. Movement-matched CS+ and CS- trial subsets were generated using the same procedure as in Supplementary Figure 3, applied to value conditions rather than cue locations. Voxel color indicates value-kernel amplitude after spatial smoothing, with red indicating CS+ preference and blue indicating CS- preference. Red contours denote the convergent value-encoding region identified in the full dataset, and blue contours denote the divergent value-encoding region identified in the full dataset. (D) Overlay of significant convergent and divergent value-encoding contours from the full dataset and movement-matched trials. Convergent value-encoding regions significantly overlapped across analyses (p < 0.0001), as did divergent value-encoding regions (p = 0.006), indicating that the spatial organization of value encoding was preserved after controlling for CS+/CS- movement differences. (E) Overlay of significant convergent and divergent value-encoding contours from the full dataset and from trials without short-latency prelicking during the cue period. Convergent value-encoding regions significantly overlapped across analyses (p < 0.0001), as did divergent value-encoding regions (p = 0.002), indicating that the spatial organization of value encoding was preserved after excluding trials with short-latency licking. Spatial overlap was assessed using the spatial overlap test described in Methods. Asterisks indicate CS+-versus CS- comparisons: *p < 0.05, **p < 0.01, ***p < 0.001; ns, not significant.

**Supplementary Figure 7.**
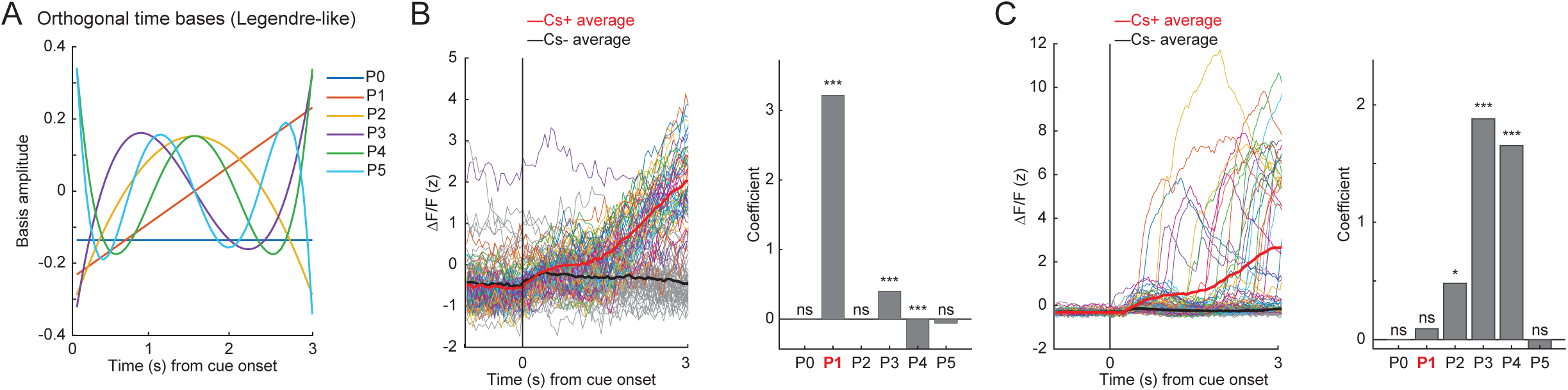
Orthogonal time-basis model for identifying monotonic ramping (A) Orthogonal temporal basis functions, P0-P5, used to model cue-period activity. (B) Example dSPN ramping site analyzed with the orthogonal time-basis model. Left, individual trial responses and CS+ and CS- trial averages aligned to cue onset. Right, fitted coefficients for each basis term. This site was dominated by the linear term, P1, consistent with monotonic ramping. (C) Example non-ramping site analyzed with the orthogonal time-basis model. Although the trial-averaged response appeared ramp-like, individual trial responses were not monotonic and the fitted response was dominated by higher-order basis terms, consistent with a non-ramping late response. Asterisks indicate significant fitted basis coefficients: *p < 0.05, **p < 0.01, ***p < 0.001; ns, not significant.

**Supplementary Figure 8.**
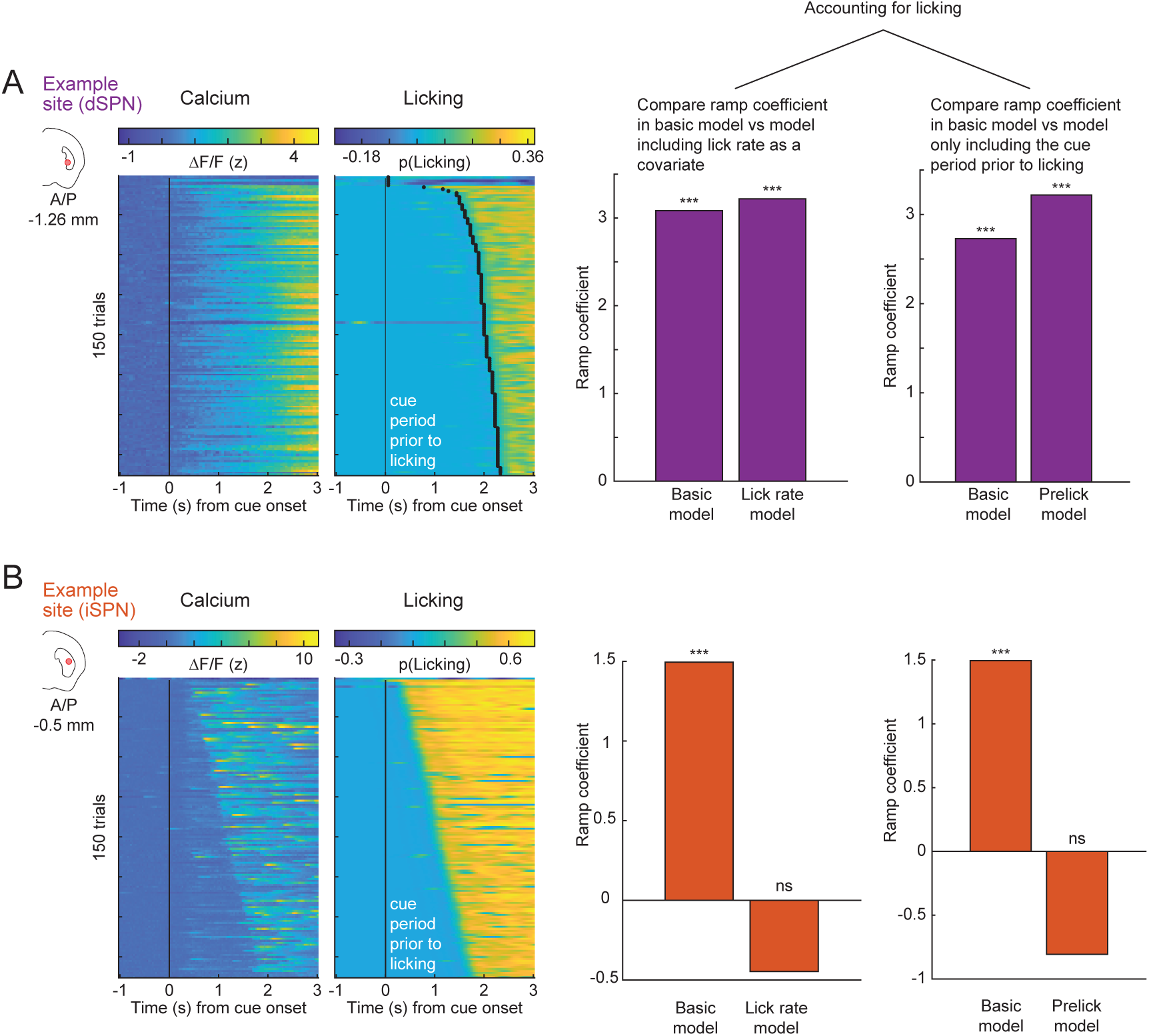
Licking controls for delay-period ramping (A) Example dSPN ramping site in which ramping persisted after licking-control analyses. Left, trial-by-trial calcium activity and licking aligned to cue onset; each row represents one trial and the black line marks first lick onset. Right, ramp coefficients from the basic orthogonal time-basis model, the model including lick rate as a covariate, and the model restricted to the cue period before lick initiation. (B) Example iSPN site in which apparent ramping was abolished by licking-control analyses. Left, trial-by-trial calcium activity and licking aligned to cue onset; each row represents one trial and the black line marks first lick onset. Right, ramp coefficients from the basic orthogonal time-basis model, the model including lick rate as a covariate, and the model restricted to the cue period before lick initiation. Asterisks indicate significant ramp coefficients: ***p < 0.001; ns, not significant.

**Supplementary Figure 9.**
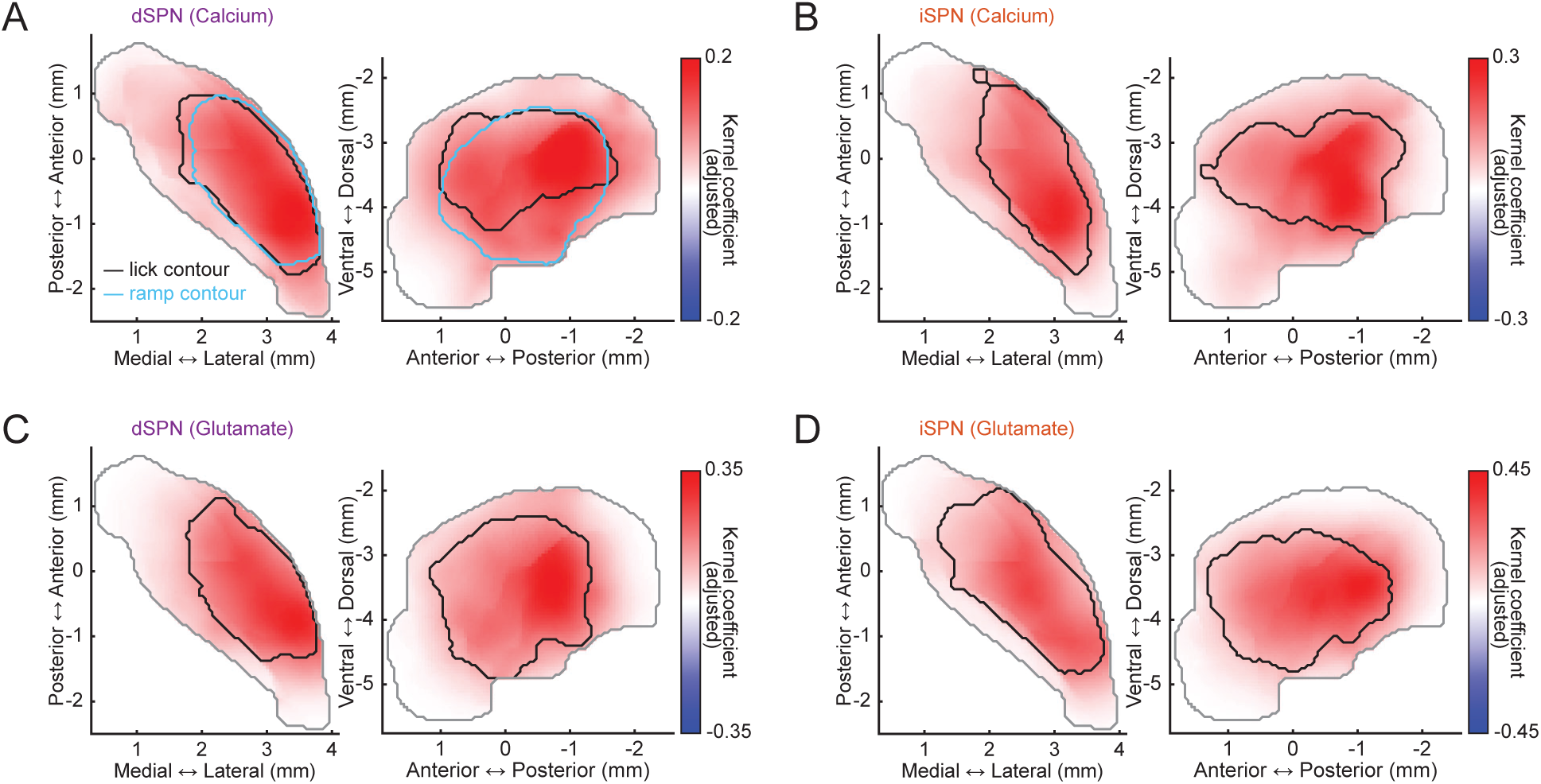
Lick-related calcium and glutamate signals in posterior ventral striatum (A) Smoothed horizontal and sagittal maps of lick-kernel strength in dSPN calcium recordings. Voxel color indicates lick-kernel coefficient amplitude after spatial smoothing. Black contours mark the significant lick-encoding region, and cyan contours mark the significant dSPN ramping region. (B) Same as (A), for iSPN calcium recordings. (C) Same as (A), for dSPN-targeted glutamate recordings. (D) Same as (A), for iSPN-targeted glutamate recordings.

**Supplementary Figure 10.**
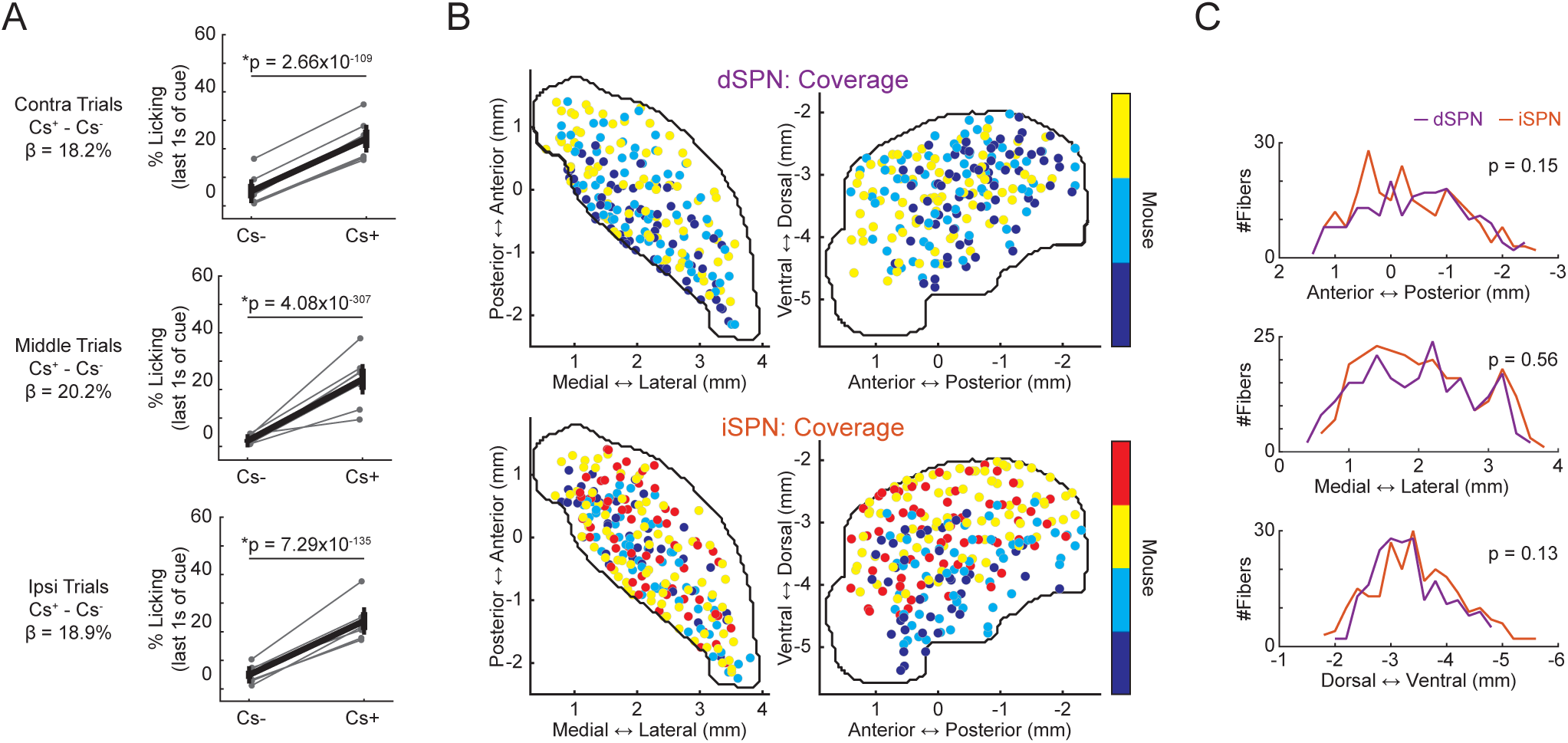
Behavior and recording coverage in glutamate-recording mice (A) Cue discrimination in the learned phase for contralateral, middle, and ipsilateral cue locations, defined relative to the implanted hemisphere, in the glutamate-recording cohort. Each line connects CS- and CS+ learned-phase values from one mouse. Anticipatory licking was quantified as lick probability during the final second before reward delivery. CS+ and CS- lick probabilities were compared using a linear mixed-effects model with cue type as a fixed effect and random intercepts for mouse and session. Statistical significance was determined from the p value of the cue type term. contralateral: β = 18.2%, p = 2.66 × 10^-109^; middle: β = 20.2%, p = 4.08 × 10^-307^; ipsilateral: β = 18.9%, p = 7.29 × 10 ^-135^. (B) Horizontal and sagittal distributions of striatal recording sites from dSPN-targeted and iSPN-targeted glutamate-imaging cohorts. Each circle indicates one fiber location, and circle color denotes the individual mouse from which that recording site was obtained. (C) Sampling coverage along the anterior-posterior, medial-lateral, and dorsal-ventral axes. Purple and orange lines indicate the number of dSPN-targeted and iSPN-targeted glutamate recording sites, respectively, at each coordinate value. Coverage along each axis was comparable between dSPN-targeted and iSPN-targeted glutamate cohorts. Two-sample Kolmogorov-Smirnov tests, anterior-posterior: p = 0.15; medial-lateral: p = 0.56; dorsal-ventral: p = 0.13. Asterisks indicate CS+ versus-CS- comparisons: *p < 0.05.

